# Internalized α-synuclein fibrils become truncated and resist degradation in neurons while glial cells rapidly degrade α-synuclein fibrils

**DOI:** 10.1101/2024.06.05.597615

**Authors:** Md. Razaul Karim, Elizabeth Tiegs, Emilie Gasparini, Riley Schlichte, Scott C. Vermilyea, Michael K. Lee

## Abstract

Parkinson’s disease (PD) and other α-synucleinopathies are characterized by the intracellular aggregates of α-synuclein (αS) believed to spread via the cell-to-cell transmission. To understand the contributions of various brain cells to the spreading of αS pathology, we examined the metabolism of αS aggregates in neuronal and glial cells. In neurons, while the full-length αS rapidly disappeared following αS PFF uptake, truncated αS accumulated with a half-life of days rather than hours. Epitope mapping and fractionation studies indicate that αS fibrils internalized by neurons was truncated at the C-terminal region and remained insoluble. In contrast, microglia and astrocytes rapidly metabolized αS fibrils as the half-lives of αS fibrils in these glial cells were <6 hours. Differential uptake and processing of αS fibrils by neurons and glia was recapitulated in vivo where injection of fluorescently labeled αS fibrils initially accumulated in glial cells followed by rapid clearance while neurons stably accumulated αS fibrils at slower rate. Immunolocalization and subcellular fractionation studies show that internalized αS PFF is initially localized to endosomes followed by lysosomes. The lysosome is largely responsible for the degradation of internalized αS PFF as the inhibition of lysosomal function leads to the stabilization of αS in all cell types. Significantly, αS PFF causes lysosomal dysfunction in neurons. In summary, we show that neurons are inefficient in metabolizing internalized αS aggregates, partially because αS aggregates cause lysosomal dysfunction, potentially generating aggregation-prone truncated αS. In contrast, glial cells may protect neurons from αS aggregates by rapidly clearing αS aggregates.

## Introduction

Neurodegenerative diseases characterized by the presence of α-synuclein (αS) aggregates are classified as α-synucleinopathies, including Parkinson’s disease (PD) and Dementia with Lewy Bodies (DLB). In PD and DLB, progressive neurodegeneration is accompanied by the presence of cytoplasmic αS aggregates called Lewy bodies (LB) and Lewy neurites (LN) [1]. αS is a highly conserved protein consisting of 140 amino acids that is predominantly expressed in neurons and enriched in presynaptic terminals. Under pathological conditions, αS adopts β-sheet conformations stabilized in oligomeric and/or fibrillar structures [2,3]. A series of studies now establish that αS pathology can spread from cell-to-cell [4–8].

While the mechanistic details of cell-to-cell spreading of αS pathology are still being uncovered, the general view is that a donor cell releases αS oligomer/aggregates, and the neighboring recipient cell internalizes the αS aggregates. Once in the recipient cell, internalized αS is thought to induce aggregation of αS in the recipient cell, possibly by acting as a seed for further aggregation. Although the details of how αS aggregates ultimately template or induce aggregation of endogenous αS are not fully understood, the metabolism/degradation of exogenous αS by recipient cells is an important factor in the spreading of αS pathology. In this regard, studies show that exogenous αS is taken up via endocytosis and trafficked via the endo-lysosomal pathway [9]. Presumably, the αS is eventually degraded via the major proteolytic pathways in the cell, including autophagy-lysosome and ubiquitin-proteasome systems [10,11].

In addition, studies show that different cell types in the brain may differentially metabolize internalized αS. For example, astrocytes can protect neurons from αS toxicity by competing for uptake of extracellular αS and degrading αS, presumably in the lysosome [6]. Similarly, microglia may coordinately degrade internalized αS via tunneling nanotubes and lysosomes [12,13]. However, these studies use cells that are exposed to relatively large amounts of αS aggregates for long periods. Thus, how different neural cell types handle αS PFF immediately following uptake is not completely understood.

To better understand the contributions of various brain cell types in the spreading of αS pathology, we examined αS trafficking and metabolism at short time points following uptake in the major brain cell types (neurons, astrocytes, microglia, and oligodendrocytes). We show that microglia and astrocytes rapidly metabolize αS PFF where the half-lives of αS PFF in these glial cells are ∼5 hours. In neurons, while the full-length αS rapidly disappears following αS PFF uptake, substantial amount of C-terminally truncated αS stably accumulates and persists for days. We also confirm that internalized αS PFF is trafficked to endosomes followed by lysosomes. In glial cells, lysosome can completely degrade internalized αS. In neurons, while αS PFF is trafficked to lysosomes but is not fully degraded. Collectively, our results show that different brain cell types differentially metabolize internalized αS PFF and that glial cells could attenuate cell-to-cell transmission of α-synucleinopathy by reducing available αS aggregates.

## Results

### Exogenous αS fibrils accumulate as a truncated species in neurons

While neurons internalize both exogenous αS monomers and fibrils, information regarding how neurons metabolize αS monomers and fibrils following the intracellular uptake is incomplete. We treated primary cultures of cortical neurons with human αS monomers and αS PFF to study the kinetics of intracellular αS metabolism. To minimize the impact of continuous uptake of exogenous αS in the culture media in the analysis of αS metabolism, we removed exogenous αS after 2h of uptake period. Briefly, the neurons were transiently exposed to exogenous αS for 2h and then washed with PBS with and without Trypsin-EDTA to remove any excess PFFs attached to the outer membrane of neurons (**Fig. S1**). The amount of residual αS following PBS wash was comparable to the cells washed with PBS containing Trypsin, which proteolyzes any residual αS attached to the outer membrane of neurons. Thus, the majority (∼90%) of the remaining αS following the PBS wash represents internalized αS as they were resistant to the trypsin wash (**Fig. S1B**).

In neurons treated with αS monomers, the transient increase in total αS level rapidly decreases to the endogenous αS levels within 3 hours (**Fig. 1A, B**). Immunoblot analysis for the exogenous human αS, using the anti-HuαS antibody [14], confirms the rapid loss of human αS monomer in neurons (**Fig. 1A**). In neurons treated with αS PFF, internalized αS persists for a much longer period, with the approximate half-life of 12 hours for the full-length αS (αS^FL^) (**Fig. 1 C, D**), indicating that the internalized αS PFF is more stable than the αS monomer. Significantly, αS PFF treatment led to the appearance of truncated αS (αS^Δ^) metabolites with a major species resolving at ∼11 kDa and corresponded to the disappearance of αS^FL^ (**Fig. 1C, D**). Significantly, αS^Δ^ stably remains even after 48 hours (**Fig. 1C, D**). Finally, αS PFF treatment leads to the stable accumulation of high molecular weight (HMW) SDS-resistant αS oligomers with major Syn-1 reactive species resolving at ∼37kDa (**Fig. 1C, lower panel**). These results show that while neurons can rapidly degrade internalized αS monomers, internalized αS aggregates remain stable for a prolonged period. We note that “pulsing” αS PFF exposure by removing uninternalized αS PFF is important for accurate determination of αS PFF metabolism as neurons in unwashed cultures continue to internalize more αS PFF over 24-hour period (**Fig. S1D**)

**Figure 1.**
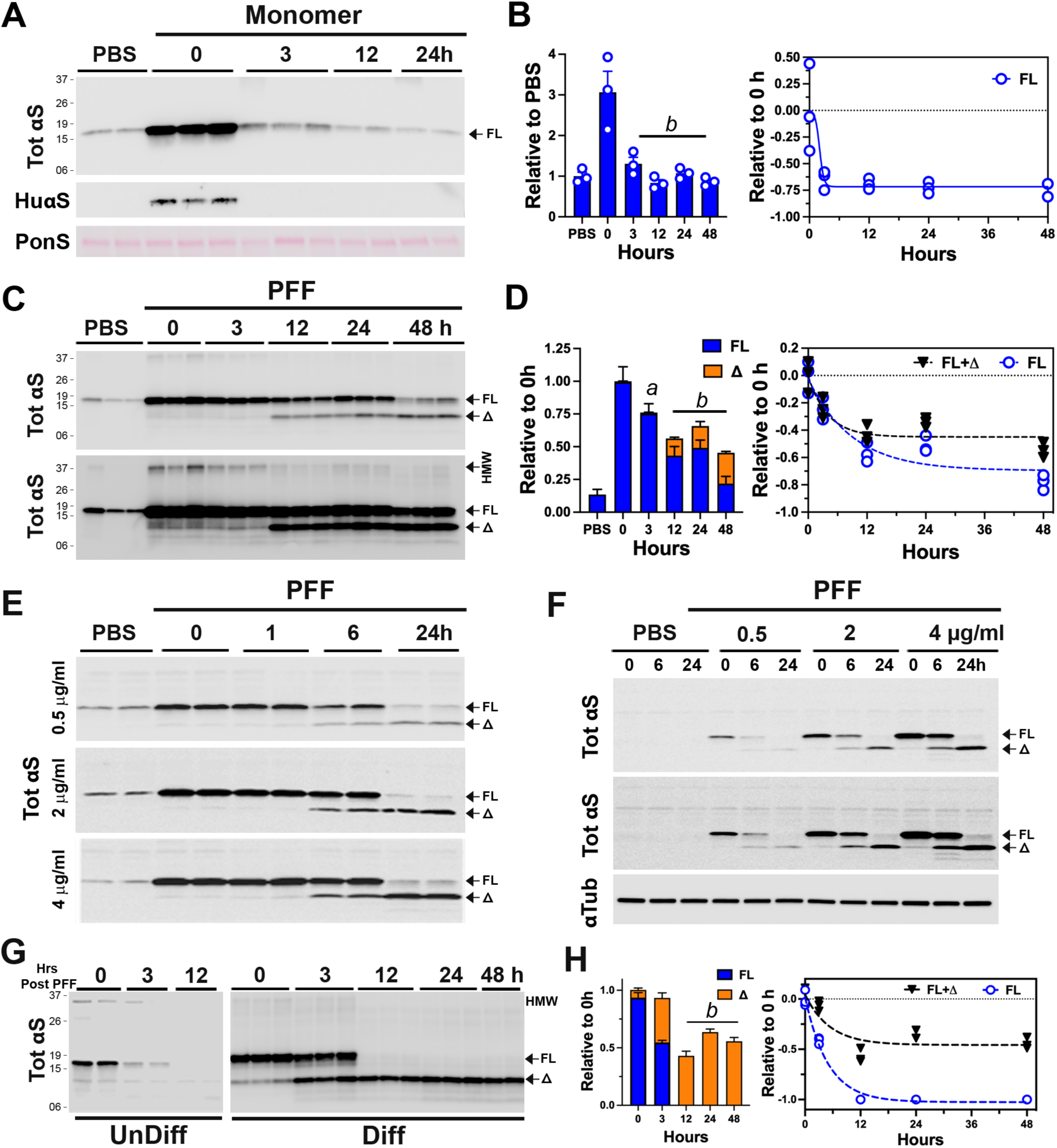
Exogenous αS pre-formed fibril stably accumulates as a truncated variant in primary cortical neurons. Primary cortical neurons (PCN) cultured from newborn C57BL6/J mouse brains at 7 days in vitro (DIV) were used. PCN were pre-incubated with 4 µg/ml (unless otherwise indicated) **(A, B)** monomeric αS or **(C-F)** αS PFF for 2h and washed to removed extracellular αS or αS PFF that was not internalized. Cells were provided with fresh media and then incubated for the indicated time before harvesting. **A)** Following αS monomer treatment, cell lysates were immunoblotted for total (Tot) αS or Hu-αS. Equal loading of proteins was verified using total protein stain (Ponceau S, PonS). **B)** Quantitative analysis of TotαS immunoblot shows that internalized αS monomers are rapidly metabolized within 3 hours of internalization. Mean±SEM; n=3. *b, p<0.001* vs 0-hour, One-way ANOVA. **C)** Immunoblot analyses for TotαS following PFF treatment show that internalized full-length (FL) αS is truncated (Δ) and stably accumulate as the truncated species. Shown are two different exposure levels to show details. **D)** Quantitative analysis of TotαS immunoblot in **(C)**. Bar and line graph show the kinetics of degradation and accumulation of αS species over time. Mean±SEM; n=3. *a, p<0.01*; *b, p<0.001* vs. total αS (FL+Δ) at 0-hour, One-way ANOVA. **E)** To determine if the amount of αS PFFs internalized impacts the truncation and/or stability of internalized αS PFFs, PCN was treated with 0.5-, 2-, and 4-μg/ml of αS PFF and the status of TotαS was analyzed at indicated times. Even at very low amounts of PFFs (0.5 μg/ml), internalized PFFs are rapidly truncated and stably accumulate. **F)** To determine if endogenous αS impacts the metabolism of internalized αS PFFs PCN established from αS KO mice were treated with αS PFFs. The results show that the lack of endogenous αS does not impact the metabolism of internalized αS PFFs. Shown are two different exposures for TotalαS to show details. Equal loading of proteins was verified by immunoblotting for α-tubulin (αTub). G) Undifferentiated (UnDiff) or neuronally differentiated (Diff) hippocampal cell line (CLU198) were treated with 4 µg/ml αS PFF and internalized αS PFF were analyzed at indicated time. In UnDiff cells, internalized αS PFF is rapidly degraded by 12 h. In Diff cells, truncated αS (Δ) stably accumulates. **H)** Quantitative analysis of immunoblot shown in **(G,** Diff). The bar graph shows the relative status of full-length (FL) and truncated (Δ) αS levels over time, confirming the stable accumulation of truncated αS. Mean±SEM; n=3. *b, p<0.001* vs total αS (FL+Δ) at 0-hour, One-way ANOVA.

Because large amounts of internalized αS PFF may overload the cellular trafficking and degradation system, we asked whether the truncation and stability of internalized αS PFF depended on the amounts of αS PFF internalized. Cultured neurons were treated with 0.5, 2, or 4 μg/ml of αS PFF and the status of internalized αS was analyzed at various time following the PFF treatment (**Fig. 1E**). The results show that even in neurons exposed to the lowest amount of αS PFF (0.5 μg/ml, ∼30 ηM), internalized αS PFF stably accumulates as truncated species with similar kinetics to neurons treated with larger amounts of αS PFF (**Fig. 1E**). Thus, generation and accumulation of truncated αS is an intrinsic property of αS PFF metabolism in neurons rather than a secondary effect of αS PFF overloading the protein degradation system. We also determined if endogenous αS expression impacts the metabolism of exogenous αS PFF by treating cortical neurons lacking αS expression, established from αS^KO^ mice, with αS PFF (**Fig. 1F**). Comparison with the wildtype (WT) mouse neuron (**Fig. 1E**) showed that αS PFF is identically metabolized in both WT and αS^KO^ neurons.

We previously showed that in neuronal cell lines and neurons, αS turnover slows with neuronal differentiation and maturation [15]. Thus, we examined whether neuronal differentiation impacts the metabolism of internalized αS PFF. For this study, we used the mouse embryonic hippocampal cell line (CLU198) that can be induced to differentiate into neuronal phenotype [16] and can efficiently internalize αS PFF (**Fig. S1C**). In undifferentiated CLU198 cells, internalized αS PFF, including the HMW species, disappears rapidly (half-life of <3 h) with no intermediate accumulation of αS^Δ^ species (**Fig. 1G**). In contrast, in neuronally differentiated CLU198 cells, the disappearance of αS^FL^ was accompanied by the stable accumulation of HMW αS and αS^Δ^ species (**Fig 1G**), even at 48h following uptake (**Fig. 1G, H**). Thus, the metabolism of αS PFF in neuronally differentiated CLU198 neuroblastoma cells was very similar to that seen in primary neurons (**Figs. 1C, D, G, H**). Analysis of αS PFF metabolism in a human neuroblastoma cell line (SH-SY5Y cells; RRID:CVCL_0019) (**Fig. S1E**) showed a similar pattern of αS metabolism as seen in CLU198 cells. Undifferentiated SH-SY5Y cells efficiently degraded αS PFF while in differentiated SH-SY5Y cells, internalized αS PFF stably accumulated as αS^Δ^ (**Fig. S1E**). Collectively, our results confirm that neuronal differentiation and maturation are associated with the distinct metabolism of αS PFF where the αS^FL^ is rapidly processed and αS stably accumulates as αS^Δ^.

### Internalized αS PFF is rapidly cleared by astrocytes and microglia but not oligodendrocytes

The differences in αS PFF metabolism between undifferentiated and differentiated neuronal states in the neuronal cell lines suggest that glial cells in the brain may be more efficient in metabolizing internalized αS PFF than neurons. Thus, we examined the clearance/metabolism of αS PFF in primary cultures of microglia, astrocytes, and oligodendrocytes. The purity of the cell types in cultures by immunohistochemical analysis for the cell type markers (**Fig. S1F-H**). As with neurons (**Fig. S1 B, C)**, analysis of cells washed with trypsin following 2h exposure to αS PFF showed all cell types internalized αS PFF (**Fig. S1F-H**).

To define the clearance of internalized αS PFF, the cell lysates were collected at different times following the wash and analyzed for internalized αS. Our results show that both microglia and astrocytes efficiently metabolized αS PFF with the approximate half-life of 6h for microglia and 3h for astrocytes (**Fig. 2A, B**). Further, no obvious accumulation of αS^Δ^ occurs in either of the cell types (**Fig. 2A, B**). Analysis of human embryonic kidney cell line, HEK293 showed that, like the other non-neuronal cells, the HEK293 cells efficiently degraded the internalized αS PFF (**Fig. S1I, J**).

**Figure 2.**
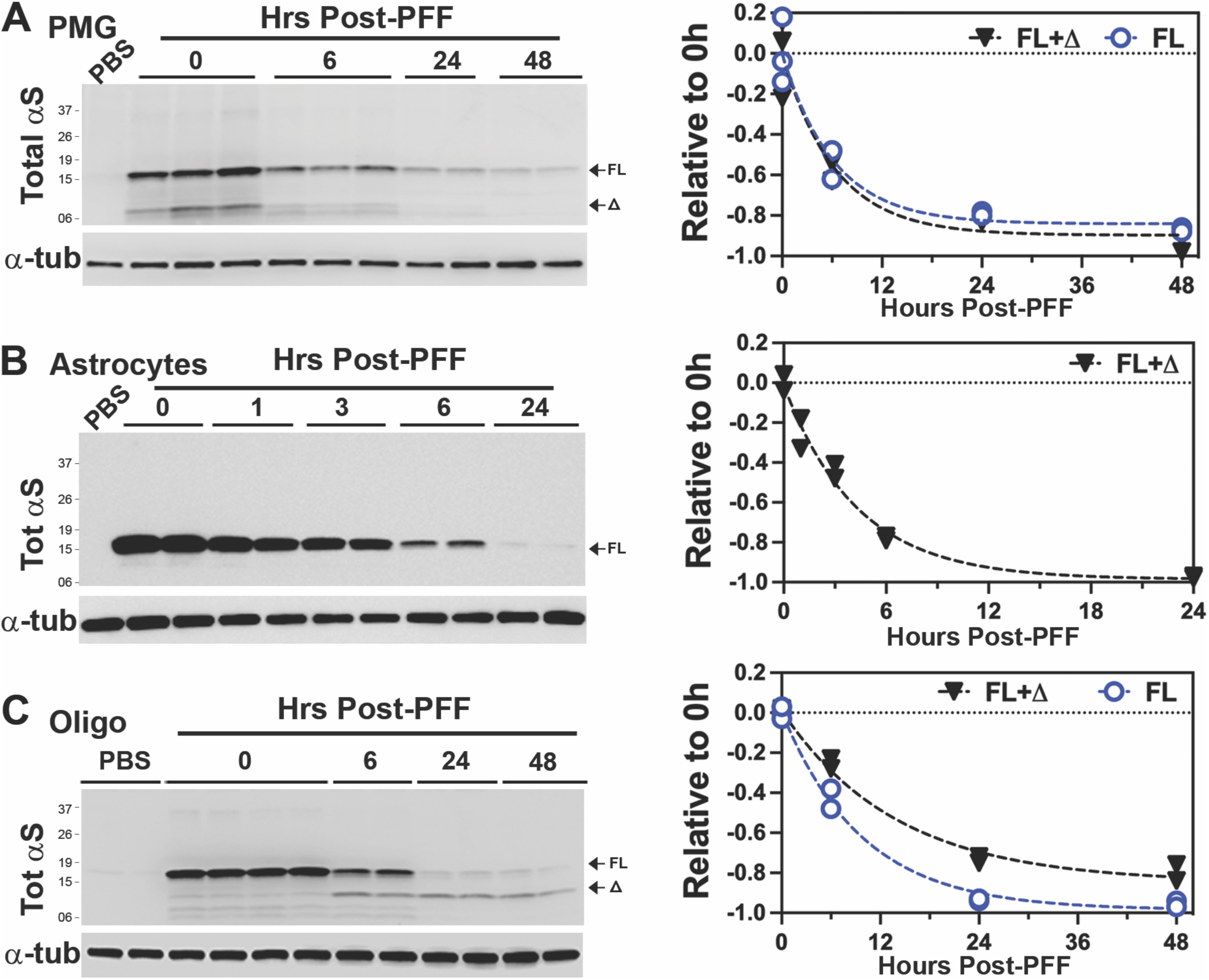
Glial cells rapidly degrade internalized αS PFF. **(A)** Primary microglia (MG), **(B)** Primary astrocyte (Astro), and **(C)** Primary oligodendrocyte (Oligo) cultures were established from newborn C57BL/6 mouse brains. Cells were pre-incubated with 4 µg/ml αS PFF, harvested at indicated times post-washing, and immunoblotted for TotαS. α-tubulin (αTub) was used as a loading control. Quantitative analysis of the immunoblots (Graphs) shows the rate of decrease in TotαS. Note that most of the internalized αS PFFs are degraded within 12-24 h with a much shorter half-life (∼6 hours) than in neurons. Further, astrocytes and microglia do not accumulate truncated αS seen in neurons. Significantly, oligodendrocytes stably accumulate truncated αS (Δ), albeit at lower levels than in neurons. Mean±SEM; n=2 per time point.

Significantly, primary oligodendrocytes show distinct αS PFF metabolism compared to the other glial cell types. While the oligodendrocytes rapidly metabolized αS^FL^, with a half-life of ∼6h, oligodendrocytes also generated αS^Δ^ species that remained stable for a prolonged period (**Fig. 2C**). Thus, the oligodendrocytes resemble neurons in the metabolism of αS PFF by generating and accumulating αS^Δ^.

Thus far, our results show that microglia and astrocytes efficiently internalized and degraded αS PFF. However, neurons and oligodendrocytes did not fully metabolize αS PFF as αS^Δ^ accumulated in these cells. While accumulation of αS^Δ^ in αS PFF-treated neurons has been reported [17], this report is first to show that non-neuronal cells do not accumulate αS^Δ^ and that oligodendrocytes also accumulate αS^Δ^ following internalization of αS PFF. It is significant that both neurons and oligodendrocytes exhibit similar metabolism of internalized αS PFF as these are the two major cell types known to develop α-synuclein pathology in brain [18–20].

### Differential accumulation of αS PFF in neural cells types occurs in vivo

We examined if the exogenous αS internalization and accumulation is different between neurons and glia in brain. We injected αS PFF labeled with Alex Fluor-488 (αS PFF-488) into the CA2/3 area of mouse hippocampus (**Fig. S2A, B**). The brains were collected at 3 hours and 24 hours post injection and processed for cellular localization of the αS PFF-488(**Fig. S2A, B**). The cellular localization of the αS PFF-488 was examined at 2 different locations, CA1 pyramidal cell layer and molecular layer (ML) dorsal to Dentate Gyrus (**Fig. S2B**). The sections were also immunostained to identify astrocytes (GFAP), microglia (Iba1) and neurons (NeuN) (**Fig. 3**).

**Figure 3.**
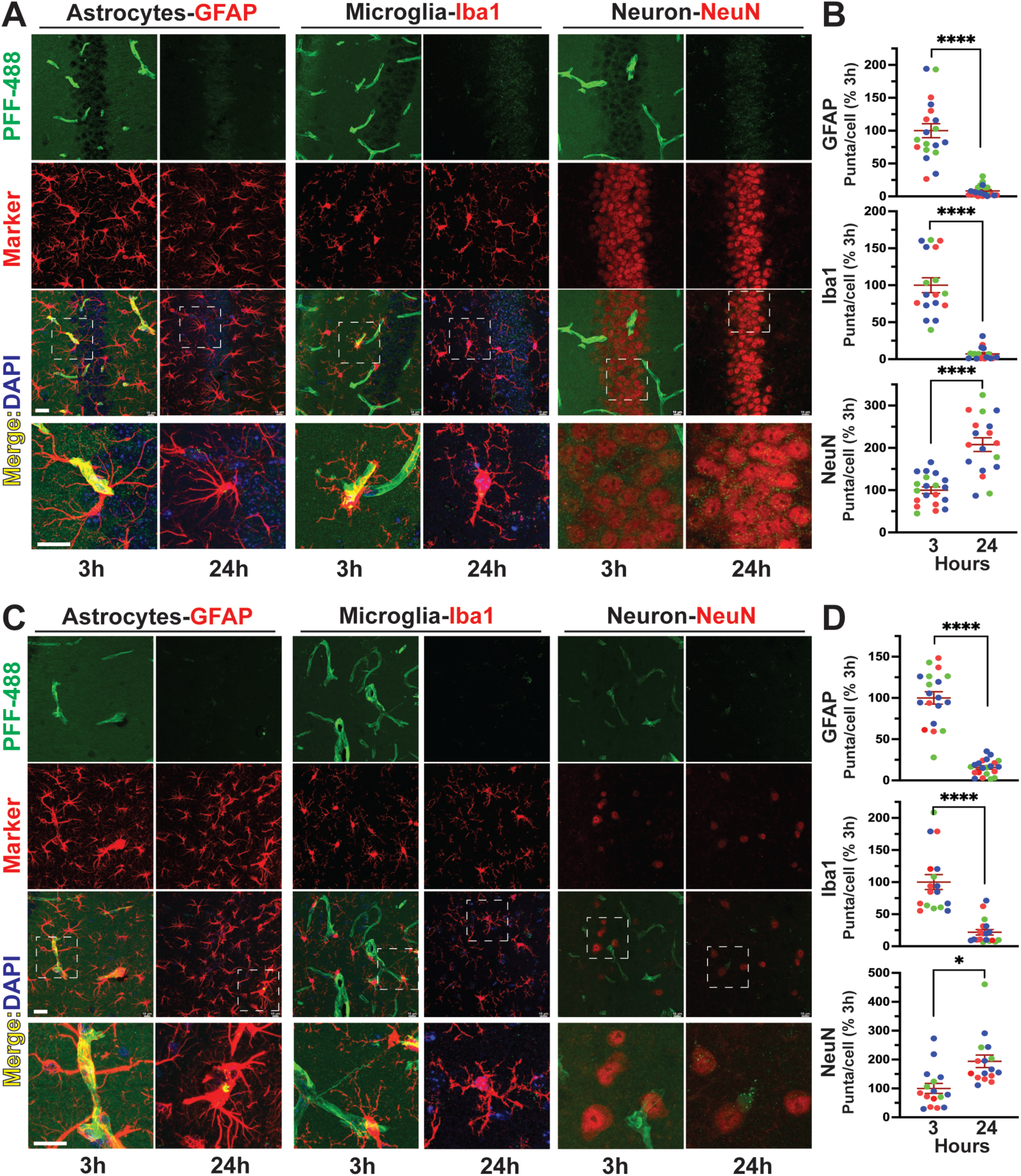
Exogenous αS PFF injected into brain is rapidly cleared by glial cells but accumulate in neurons. AF-488 labelled αS PFF was streotaxically injected into mouse brain (hippocampus/cortex) and the localization of AF-488 was evaluated in brains harvested at 3h and 24h post injection. 20 µm thick free-floating section was immunostained with various cellular markers: Astrocytes, GFAP; Microglia, Iba1; Neuron, NeuN for CA1 (A-B) and Molecular Layer (ML)-dentate gyrus region (C-D) of Hippocampus. Multiple confocal images was used to count the specific cellular colocalization of PFF using HALO Quantitative Image Analyzer. The graphs are plotted as percentage (%) of 3h (B,D). \*\*\*\**p*<0.0001, \**p*<0.05, unpaired *t-*test, Mean±SEM; Each value represents an average value from a single section with the sections from same animals are plotted with same color. Bar=20μm.

At 3 hours post-injection, we observe prominent accumulations of αS PFF-488 along what appears to be vesicular structures often lined with GFAP (**Fig. 3A, C**). The vascular accumulation is selective for the injected side as the αS PFF-488 signal is absent in the contra-lateral hippocampus (**Fig. S2C**). Analysis of confocal slices indicate that the some of the vascular accumulation of αS PFF-488 occurs within GFAP processes suggest that the αS PFF-488 might be concentrating along the glial end-feet lining the glymphatic space (**Fig. S2D**). Significantly, the vascular accumulation of αS PFF-488 is disappears by 24 hours post-injection (**Fig. 3A, C**). In addition to the prominent vascular accumulation, we observe colocalization of punctate αS PFF-488 signals with glial cell markers (GFAP+ or Iba1+) at 3 hours. This punctate αS PFF-488 within the glial cells are reduced at 24 hours post injection (**Fig. 3A-D**). We also observe a general increase in green-fluorescence signal throughout the neuropil in the PFF injected side, but not in the contra-side, at 3 hours that disappears by 24 hours. We propose that this represents general spread of the injected PFF through the neuropil that is cleared by 24 hours post injection (**Figs. 3, S2**). In neurons, we observed a low level of αS PFF-488 colocalizing within the NeuN+ cells at 3 hours post injection which increases significantly at 24 hours post injection (**Fig. 3 A-D**). These results indicate that glial cells, particularly astrocytes, rapidly internalize exogenous αS fibrils and the internalized αS fibrils are efficiently cleared by the glial cells within 24 hours. In neurons, while the accumulation of αS fibrils seem to occur slower, internalized αS fibrils persist past 24 hours.

### Internalized αS PFF and truncated αS remain detergent insoluble in neuronal cells

When αS PFF are sonicated to facilitate uptake by the cells, sonication leads to a significant fraction of αS PFF partitioning into buffer soluble fractions that can induce seeding of αS aggregates [21]. Our analysis also confirmed that sonicated αS PFF used to induce αS pathology contained a significant amount of soluble αS (data not shown). Thus, we asked if the internalized αS are soluble or insoluble as they are metabolized by neurons. First, we compared the solubility of monomeric αS and αS PFF internalized by differentiated CLU198 cells (**Fig. S3A, B**). CLU198 cells were treated with αS monomer (**Fig. S3A**) or αS PFFs (**Fig. S3B**) for 2 h, washed, and the cells were collected at ‘0h’ and ‘24h’. The collected samples were solubilized in triton X-100 (TX-100) and the detergent soluble and insoluble fractions were obtained by centrifugation at 100,000xg. In the monomer-treated cells, the majority of αS remained soluble at 0h and degraded by 24 h, albeit a small fraction of αS monomer, likely endogenous αS, was found in the insoluble fraction at 24 h. In the αS PFF-treated cells, the majority of αS species partitioned to the TX-100 insoluble fraction, even at 24 h. Significantly, TX-100 insoluble SDS-resistant αS oligomers resolving ∼37 kDa at 0h (**Fig. S3B, \***) resolved at slightly lower MW at 24h (**Fig. S3B, \***Δ), consistent with the truncation of the HMW αS species.

Analysis of soluble and insoluble αS in the αS PFF-treated PCN during several days following PFF treatment show that the internalized αS remained insoluble at 0 day, 3 days, and 7 days following initial αS PFF internalization (**Fig. S3C**). Consistent with very slow turnover of internalized αS PFF in neurons, ∼50% of αS remains even at 7 days post internalization.

### Neurons accumulate C-terminally truncated αS PFF

To more accurately determine the size of the αS^Δ^, we resolved αS PFF treated neuronal lysates next to the αS protein truncated at 110, 120, 130, and FL (**Fig. S4A**). We observed that the major αS^Δ^ species derived from αS PFF resolves between αS110 and αS120 with the estimated mass of ∼11.5 kDa (**Fig. S4B**).

We previously showed that αS is normally truncated at C-terminus and that C-terminally truncated αS is enriched in αS aggregates in vivo [14]. To determine if internalized αS PFF is C-terminally truncated in neurons, we performed immunoblot analysis of the soluble and insoluble fractions from PFF-treated CLU198 cells using the anti-HuαS antibody [14] that selectively recognize the HuαS C-terminal epitope (amino acids 115-122) (**Fig. 4A**). Our results show that HuαSyn antibody recognized both HMW and full length αS (αS^FL^) at 0h but the HuαS immunoreactivity was virtually absent at 24 h (**Fig 4B**). We also analyzed αS PFF-treated PCN at various times and compared the αS variants recognized by Syn-1 and anti-HuSyn antibody (**Fig. S3D**). As with the CLU198 cells, HuαSyn reactive bands disappeared with the loss of αS^FL^, indicating that the 11.5 kDa αS^Δ^ species were missing the C-terminal portion of the protein. Taken together with immunoblot analysis of the αS species detected by Syn-1 antibody (**Figs. 1, S3**), we conclude that the majority of internalized αS PFF was C-terminally truncated at 24-hours.

**Figure 4.**
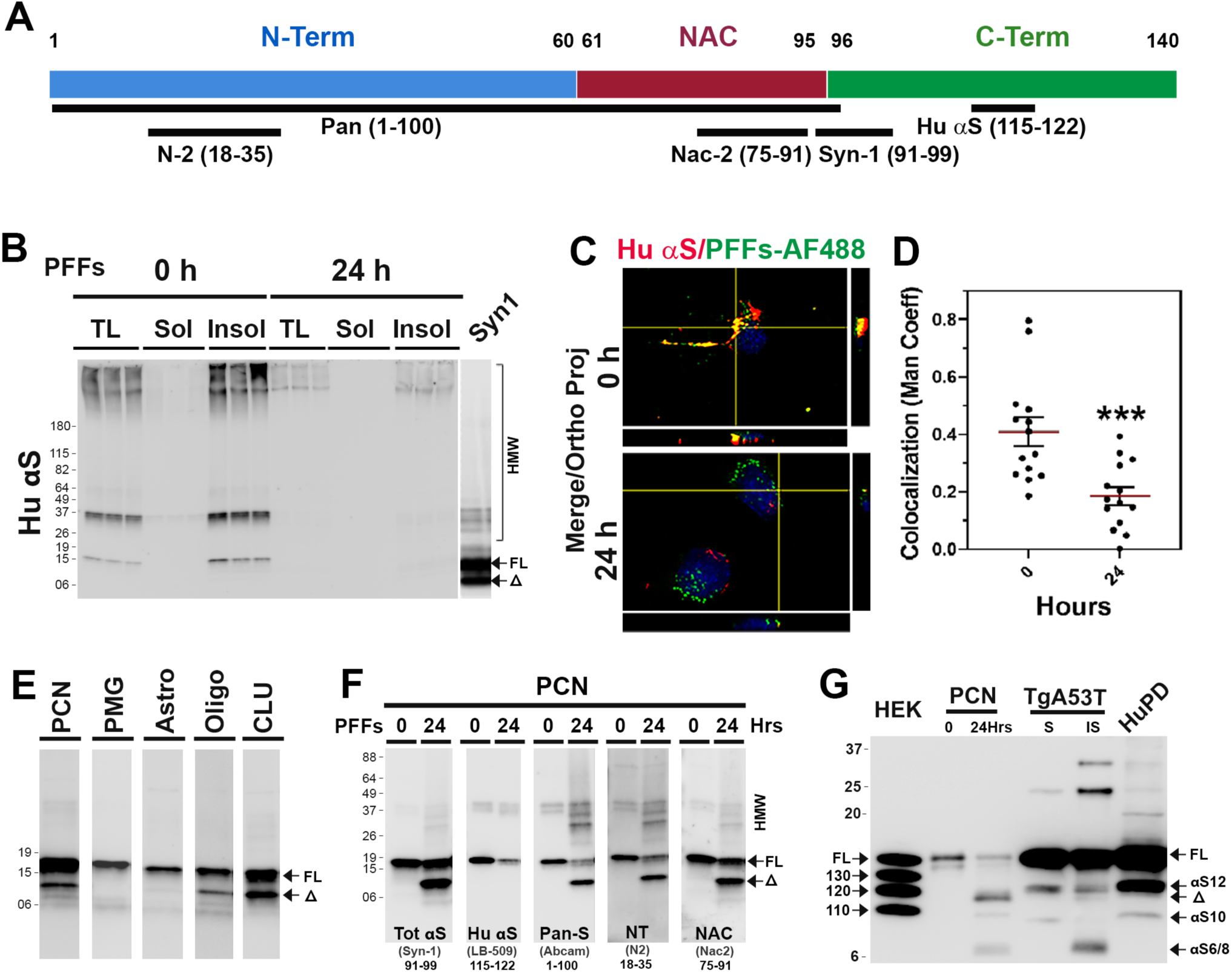
Neurons accumulate C-terminally truncated αS PFF. **(A)** Schematic representation of αS with the locations of epitopes for the anti-αS antibodies used to map the truncated αS. **(B)** TX-100 soluble (Sol) and insoluble (Insol) fractions from PFFs treated CLU198 cells, shown in Figure 4B, were immunoblotted using HuαS antibody. Note that HuαS antibody recognizes αS FL but not the truncated αS (αSΔ). **(C)** Neuronally differentiated CLU198 cells were treated for 2h with Alex Fluor 488 (PFFs-AF 488) labelled PFFs and washed to remove any extracellular PFFs-AF-488. Cells were provided with fresh media and then incubated for 0h or 24h prior to fixing and immunostained with HuαS antibody. Double immunofluorescence microscopy was used to visualize HuαS (red) and PFFs-AF 488 (green). **(D)** Co-localization (yellow) between Hu αS and PFFs-AF 488 were quantified (n=15-18 cells) by using the Manders’ coefficient (ImageJ software) and plotted. *\*\*\*p<0.001*, Unpaired *t*-test. **(E)** PFFs treated PCN, primary microglia (PMG), primary astrocytes (Astro), oligodendrocytes (Oligo), and neuronal differentiated CLU198 cells (CLU) were analyzed for αS variants. All the samples were resolved on same gel but separated and reordered for clarity. Only PCN, Oligo, and CLU accumulate αS10. (**F**) Triton X-100 (TX-100) insoluble fractions from PCN were immunoblot analyzed for Tot αS (SYN-1, epitope 91-99), C-terminal HuαS (epitope 115-122), N-terminal Pan-S (epitope 1-100), N-terminal NT-αS (epitope 18-35) and NAC-domain containing NAC-2αS (epitope 75-91). αSΔ is missing the C-terminal HuαS epitope but retains the N-terminal epitopes. (**G**) Comparison of αS variants in αS PFF-treated PCN, Human PD case (HuPD) and TgA53T mice with αS pathology. In HuPD and TgA53T mice, αS^12^ is the major truncated αS variant and αSΔ is a very minor component. In PCN, αSΔ is the dominant variant produced from internalized αS PFF.

To document the C-terminal truncation at the cellular level, neuronally differentiated CLU198 cells were treated with αS PFF labeled with Alex Fluor-488 (AF-488, green) and immunostained for HuαS and visualized using AF-647 conjugated secondary antibody, at 0 h and 24 h following αS PFF treatment (**Fig. 4C, D**). Colocalization of AF-488 (Green) with HuαSyn immunoreactivity (Red) showed high levels of colocalization between AF-488 and HuαS at 0 h. However, at 24 h following αS PFF treatment, colocalization of AF-488 with HuαS dramatically decreased, indicating that the C-terminal portion of the αS was missing from the internalized PFF at 24 h (**Fig. 4C, D**).

To further define the nature of the αS truncation, we used a variety of αS antibodies with the defined epitopes (**Fig. 4A**) to map the primary structure of αS species generated by the cells following αS PFF internalization. Immunoblot analysis with Syn-1 antibody (amino acids 91-99) [14,22] (**Fig. 4E**) showed that in PCN, CLU198, and oligodendrocytes, αS PFF treatment resulted in a major 11.5 kDa truncated αS^Δ^, while the accumulation of truncated αS species were not obvious in PMG and astrocytes (**Fig. 4E**). While small amount of αS^Δ^ was occasionally seen in PMG, these species were transient as they disappeared with the αS^FL^ (**Fig. 2A**).

Because αS aggregates in vivo contain both N- and C-terminally truncated αS [14], we used additional anti-αS antibodies with defined epitopes located at the N-terminal region, C-terminal region, and central-NAC regions (**Fig. 4A**) to map the αS^Δ^ from the αS PFF-treated PCN (**Fig. 4F**). The results show that the αS^FL^ reacts with all antibodies. The major αS^Δ^ reacted to antibodies recognizing the N-terminal (N2, pan-Syn) and the central (NAC, Syn-1) epitopes but not to the antibodies that bind to the C-terminal epitopes (HuαS, LB509). The minor truncated variants at ∼6-8 kDa only reacted to antibodies to central epitopes (Syn-1, NAC). Thus, internalized PFF was first truncated at the C-terminal region to generate ∼11.5 kDa αS^Δ^ and further truncated to remove the N-terminal region (αS^6/8^). Epitope mapping of αS species in primary oligodendrocytes (**Fig. S4C**) and hippocampal CLU198 cell lines (**Fig. S4D**) also show that these cells accumulated αS^Δ^ where C-terminal epitope was missing while retaining the N-terminal and NAC regions. In PMG, a minor truncated species at ∼6 kDa (αS^6^), missing both N- and C-terminal region is observed (**Fig. S4E**). Our results show that the stable accumulation of C-terminally truncated αS is a common feature of neuronal cells (PCN and CLU198 cells) and the oligodendrocytes, cell types that are associated with α-synucleinopathies.

To determine if the αS^Δ^ seen in PFF treated neurons accumulate *in vivo* with αS pathology and to determine how the αS^Δ^ compares to previously identified C-terminally truncated αS [14], we compared the αS from PFF treated neurons with the insoluble αS from the transgenic mouse expressing A53T mutant human αS (TgA53T) and human brain affected by α-synucleinopathy (**Fig. 4G**). Insoluble aggregate recovered from TgA53T is qualitatively similar to the aggregates recovered from human PD cases [14] (Fig. 4G) with major C-terminally truncated species at ∼12 kDa (αS^12^). Comparison of TgA53T and human PD lysates with αS PFF treated neurons show that αS11.5 resulting from internalized αS PFF is qualitatively different from the insoluble αS aggregates recovered from the TgA53T mouse and human PD case. Specifically, the major truncated species in the TgA53T mouse model and in human PD case resolve at ∼12 kDa (αS^12^) [14], while with the αS PFF-treated neurons, αS^12^ was not seen and 11.5 kDa αS^Δ^ was most abundant (Fig. 4G). We previously showed that insoluble αS from TgA53T model and human PD case contains a major C-terminally truncated species at ∼12 kDa (αS^12^), a minor C-terminal species 10 kDa species (αS^10^), and another truncated species at ∼6-8 kDa lacking both N- and C-terminal regions (αS^8^) [14]. Thus, our comparative analysis shows that 11.5 kDa C-terminally truncated αS is an unique product of αS PFF metabolism by neurons.

### Internalized αS PFF is trafficked to lysosomes via endosomes

Internalization of αS PFF occurs via the endosomal pathway and is trafficked to lysosomes [4,23–25]. Further, lysosomal proteases, such as cathepsins and asparagine endopeptidase (AEP), are implicated in the C-terminal truncation of αS [26–28]. Thus, we determined the time course of intracellular trafficking of αS PFF immediately after the internalization by treating the cells with αS PFF labeled with Alex Fluor-488 or −647 (AF-488 or AF-647).

To confirm that internalized αS PFF in neurons are initially trafficked to endosomes, we colocalized internalized PFF-AF488 with an early endosome marker, EEA1, at different times following PFF-AF488 treatment (**Fig. 5A-C; Fig. S5A**). At 0.5h post αS PFF treatment, a significant increase in the amount of EEA1 staining was seen compared to the levels of EEA1 staining at other times (**Fig. 5A, B; Fig. S5A**), indicating that internalization of αS PFF induced an increase in early endosomes. A strong co-localization of PFF-AF488 with EEA1 was seen at 0.5 h and the colocalization decreased at 1 h and 3 h post αS PFF treatment (**Fig. 5A, C; Fig. S5A**). Following transit through endosomes, we hypothesize that a decrease in endosomal αS represents the trafficking of internalized αS PFF to lysosomes. To confirm this, we first treated PCN with the Lysotracker Red-50 followed by transient exposure to PFF-AF488. Colocalization of the PFF-AF488 with the Lysotracker was examined at 0, 0.5, 1.0, and 3.0 h post-PFF treatment (**Fig. 5D, E; Fig. S5B**). Our results show that as the PFF-AF488 exits the endosomal compartment, PFF-AF488 accumulates in the lysosomes labeled by the Lysotracker (**Fig. 5D, E; Fig. S5B**). Analysis of PMG and astrocytes (**Fig. S5C, D**), show that internalized AF488-PFF initially co-localizes with the endosome markers followed by the lysosome markers. We also performed subcellular fractionation to obtain Lamp1-enriched lysosomal fraction and cytosolic fraction from PFF-treated CLU198 cells (**Fig. 5F**). Immunoblot analysis for αS showed that most of the αS, both αS^FL^ and αS^Δ^, is recovered with the lysosomal fractions.

**Figure 5.**
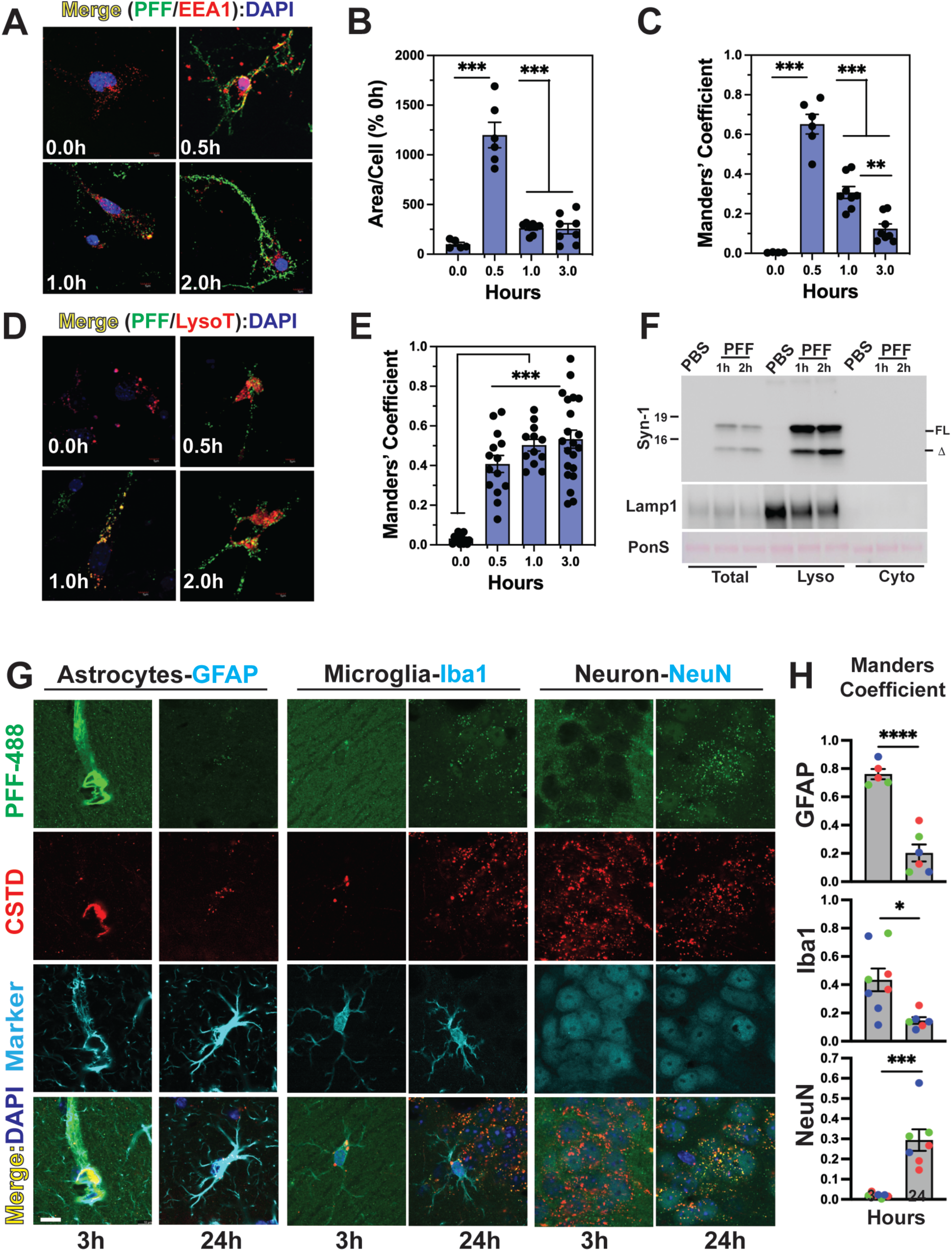
αS PFF is internalized via endosomes and trafficked to lysosomes. PCN was treated with PFF-AF488 and fixed at 0-, 0.5-, 1- and 3-h following PFF-AF488 addition. (**A**) Cells were immunostained for early endosome marker EEA1. Both EEA1 (Red) and αS PFF-AF488 (Green) were imaged by double immunofluorescence microscopy. (**B**) Quantitative analysis of the area/cell covered by EEA1 shows transient increase in early endosomes immediately following αS PFF uptake. (**C**) Colocalization of EEA1 with PFF-AF488, expressed as the Manders’ coefficient, show initial colocalization of EEA1 and PFF-AF488 followed by a progressive decrease in colocalization. (**D**) PCN treated with PFF-AF488 were labeled with Lysotracker Red (LysoT). Both PFF-AF488 (Green) and LysoT (Red) were imaged using double immunofluorescence microscopy. (**E**) Colocalization of PFFs-AF488 with LysoT, expressed as the Manders’ coefficient, shows that PFFs-AF488 is trafficked to lysosomes and continues to accumulate with lysosomes over time. (**F**) Differentiated CLU-198 cells were treated with PFFs for indicated time and cytosolic and lysosome-enriched fractions were obtained. Immunoblot analysis of the fractions shows that both αS-FL and αSΔ partitions with the lysosome fraction. The fractions were also immunoblotted for Lamp1, a lysosomal marker. *\*\*p<0.01, ***p<0.001*, One-way ANOVA. **G) Lysosomal co-localization of internalized** α**S PFF in vivo.** Brain sections from mice injected with αS PFF-488 (Fig. 3) was used to colocalize αS PFF-488 (green) with lysosomal marker cathepsin-D (CSTD/red) and cell-type marker (Astrocytes, GFAP; Microglia, Iba1; Neuron, NeuN). Co-localization of cell-specific PFF-488 with CSTD expressed as the Manders’ coefficient. Consistent with the internalization of pattern seen in Fig. 3, αS PFF-488 colocalization with CSTD significantly increased in hippocampal neuron from 3 hours to 24 hours. In astrocytes and microglia higher PFF-488 load appears at 3h but significantly decreases at 24h post injection. \**p<0.05;* \*\**\*p<0.001;* \*\*\**\*p<0.0001*, *t*-test, Mean±SEM; Each value represents an average value from a single section with the sections from same animals are plotted with same color. Bar=10μm.

Analysis of PFF-AF488 with other subcellular markers in PCN show some co-localization of αS PFF-AF488 with Lamp-2, p62, and endoplasmic reticulum marker, Grp78 (**Fig. S5E**). No significant colocalization of internalized PFF-AF488 are seen with the markers of autophagosomes (LC3) or Golgi (GM-130) (**Fig. S5E**).

To confirm that internalized αS PFF is also targeted to lysosomes in vivo, we imaged brain sections from the αS PFF-488 injected mice for colocalization of αS PFF-488 with lysosome (Cathepsin D, CSTD) (**Fig. 5G**). We observed that in astrocytes and microglia, αS PFF-488 colocalizes with CSTD at 3 hours post injection. In neurons, the low amount of αS PFF colocalizes with CSTD at 3 hours post injections but the overall number of αS PFF-488 puncta, as well as αS PFF-488 puncta colocalizing with CSTD increases at 24 hours post injection. These results confirm that in vivo, exogenous αS fibrils are rapidly internalized glial cells and degraded by the lysosomes while the neurons exhibit a slower internalization of αS fibrils that accumulate within the lysosomes.

Collectively, our results support the scenario where internalized αS PFF traffic to lysosomes via the endosomes. Further, we proposed that the truncation of αS PFF (neurons) and/or degradation of αS PFF (all cells) occurs in lysosomes. In neuronal cells, internalized αS PFFs stably accumulated in the lysosomes (**Figs. 5, S5**) and the time course of αS PFF colocalization of lysosomes mirrors the timer course of αS^Δ^ accumulation (see **Fig. 1C and E**).

### Lysosome is the major degradation machinery of αS PFF

Lysosome is an acidic organelle containing hydrolytic enzymes that are responsible for the degradation of protein aggregates, non-functioning intracellular organelles, and foreign influx [10,29]. Given that the lysosome is the predominant destination of internalized αS PFF, we hypothesize that the lysosome is responsible for the rapid metabolism of αS PFF in non-neuronal cells. To directly test the role of lysosomes in αS PFF metabolism, we used bafilomycin A1 (Baf), which inhibits v-ATPase responsible for the acidification of lysosomes. This lysosomal acidification requires the activation of resident hydrolytic enzymes to stay inside the lysosome [30].

Baf inhibition of lysosome in αS PFF-treated PCN resulted in stabilization of both αS^FL^ and αS^Δ^, particularly at 12 h post-PFF treatment (**Fig. 6A, B**). In CLU-198 cells, Baf also caused αS stabilization as in the PCN (**Fig. 6C, D**). Because CLU198 cells exhibit a faster rate of αS PFF truncations, increased levels of both αS^FL^ and αS^Δ^ with Baf treatment were obvious at 3 h following αS PFF uptake. Analysis of Triton X-100 soluble and insoluble fractions showed that the Baf treatment increased αS in the insoluble fraction (**Fig. S6A**). Further, both monomeric αS and HMW αS were stabilized by Baf treatment (**Fig. S6A**).

**Figure 6.**
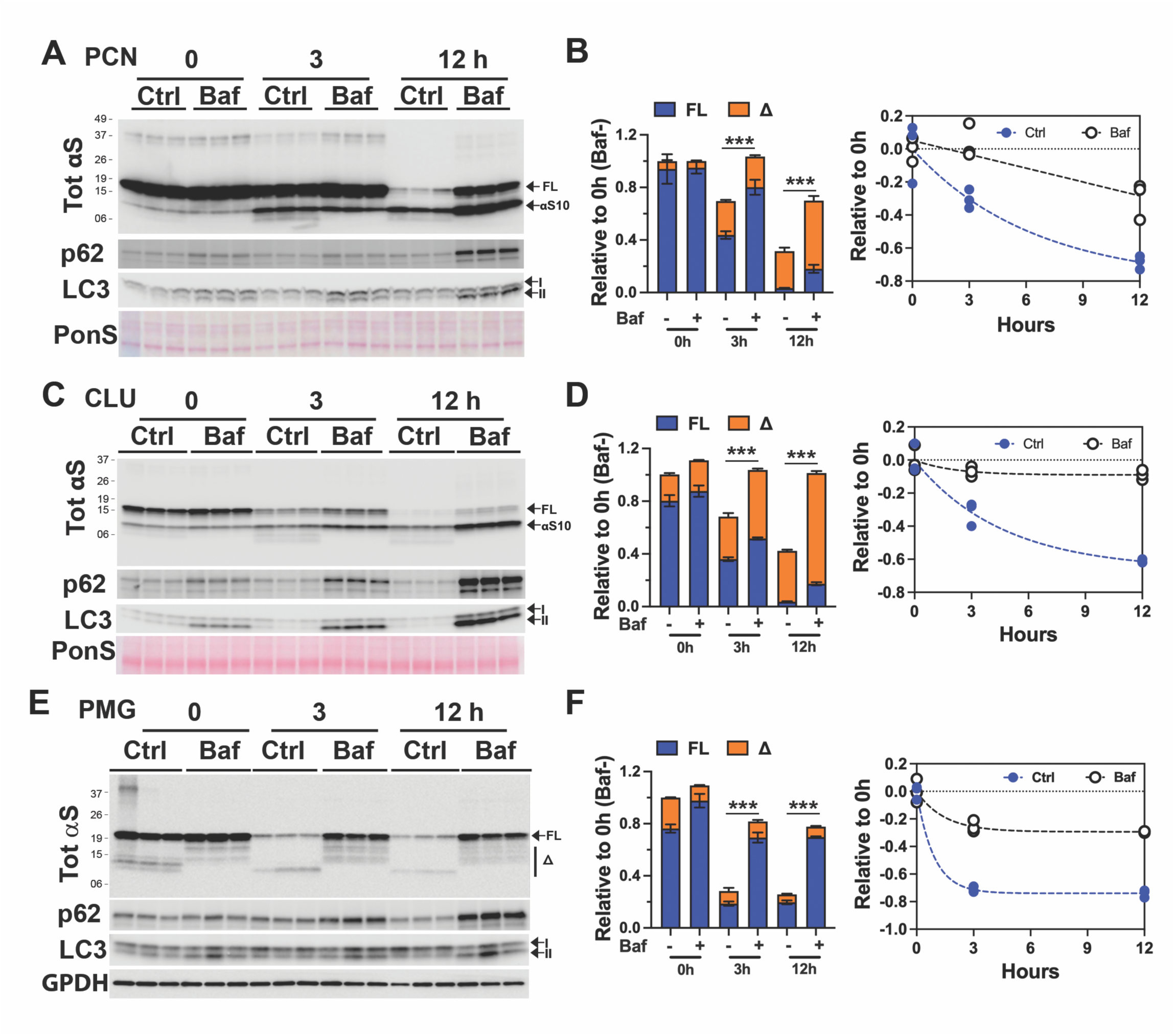
Lysosome function is required for the degradation of αS PFF in neuron and glial cells. Primary neuronal culture (PCN) **(A, B)**, neuronally differentiated CLU198 cells **(C, D)**, and Primary Microglia (PMG) **(E, F)** were pretreated with a lysosomal inhibitor bafilomycin A1 (100 ηM, Baf) or DMSO for 4h. During the last 2 h of Baf treatment, 4 µg/ml of αS PFF was added to the cells. The cells were washed and provided with fresh media containing DMSO (control, Ctrl) or Baf (100 ηM) and harvested at 0-, 3- and 12-h of incubation in fresh media. The levels of TotαS at each time point were determined by immunoblot analysis. Ponceau S (PonS) or αTub was used as loading controls. Bar graphs show that levels of total αS (FL+Δ) in Baf-treated cells are significantly higher than in controls. Further, the individual levels of FL and truncated αS (αS^10^, Δ) are significantly higher in Baf-treated cells at 12 h compared to the corresponding controls (*p<0.001*, Two-way ANOVA). The line graph and corresponding regression analysis show that Baf treatment significantly slows rate of αS degradation. Mean±SEM; n=3. *\*\*\*p<0.001*, total αS(FL+Δ) in Baf-vs Baf+, Two-way ANOVA.

We also treated non-neuronal cells with Baf to test if lysosome function is responsible for the effective degradation of internalized αS PFF. Results from PMG (**Fig. 6E, F**) show that Baf treatment leads to robust stabilization of αS PFF. Similarly, Baf treatment also prevents degradation of αS PFFs in primary astrocytes (**Fig. S6B**) and HEK293 cells (**Fig. S6C**).

These results show that in both neuronal and non-neuronal cells, lysosomal degradation is the predominant mode of degradation for internalized αS PFFs. Significantly, in neurons, inhibition of lysosomes by Baf does not inhibit αS truncation. Thus, while the truncated αS PFF accumulates in lysosomes, the truncation of αS PFF is independent of αS degradation.

Since the lysosome is largely responsible for the metabolism of internalized αS PFF in cells, we examined whether differences in the lysosomal content of the cells could linked to differential metabolism of internalized αS PFF in the various cell types. When the cell types used here were analyzed for the abundance of lysosome markers (Lamp-1 and Cathepsin D) (**Fig. S6D, E)**, microglia and astrocytes exhibit higher levels of lysosomal markers than PCN, indicating that the glial cells have higher lysosomal capacity than in PCN. Moreover, undifferentiated CLU198 cells (CLU-UD; **Fig. S6D, E,**) contain higher levels of lysosomal markers than the neuronally differentiated CLU198 cells (CLU-Diff; **Fig. S6D, E**,). Query of single nuclei/cell RNAseq data bases (DropViz.org; brainrnaseq.org) also show that in adult mouse and human brain, Lamp-1 and Lamp-2 expression, representing lysosome content, is 2-3 fold higher in astrocytes/microglia then in neurons (**Fig. S6F**). These results support the view that differences in the lysosomal activity contribute to the differential metabolism of internalized αS PFF in neural cell types.

Accumulation of αS fibrils in the lysosome of neuronal cells is linked to dysfunctional lysosomes [4,10]. Thus, we tested if the accumulation of truncated αS PFF in lysosomes at early time points following PFF uptake was associated with lysosomal dysfunction. We focused on neuronal cells as the rapid degradation of αS PFF by glial cells excludes the possibility of lysosomal dysfunction by αS PFF in glial cells. Differentiated CLU198 cells were treated with saline or αS PFF-AF488 and at 6- and 24-hours post-αS PFF uptake, the lysosomal function was evaluated on live cells using the Magic Red Cathepsin B activity kit (**Fig. 7A, Fig. S7A**). Our results show that both 6- and 24-hours post-PFF uptake, the levels of Cathepsin B activity were notably lower in α PFF-treated cells (**Fig. 7A, Fig. S7A**). Quantitative analysis of Magic Red Cathepsin B signal at 24 hours following PFF-uptake showed that Cathepsin B function, as indicated by the area occupied by the Magic Red signal per cell, was significantly lower with αS PFF treatment (**Fig. 7B**). Immunoblot analysis for Cathepsin D in PCN and CLU cells treated with αS PFF show that αS PFF treatment leads to reduced levels of active Cathepsin D relative to pro-Cathepsin D (**Fig. S7B, C**).

**Figure 7.**
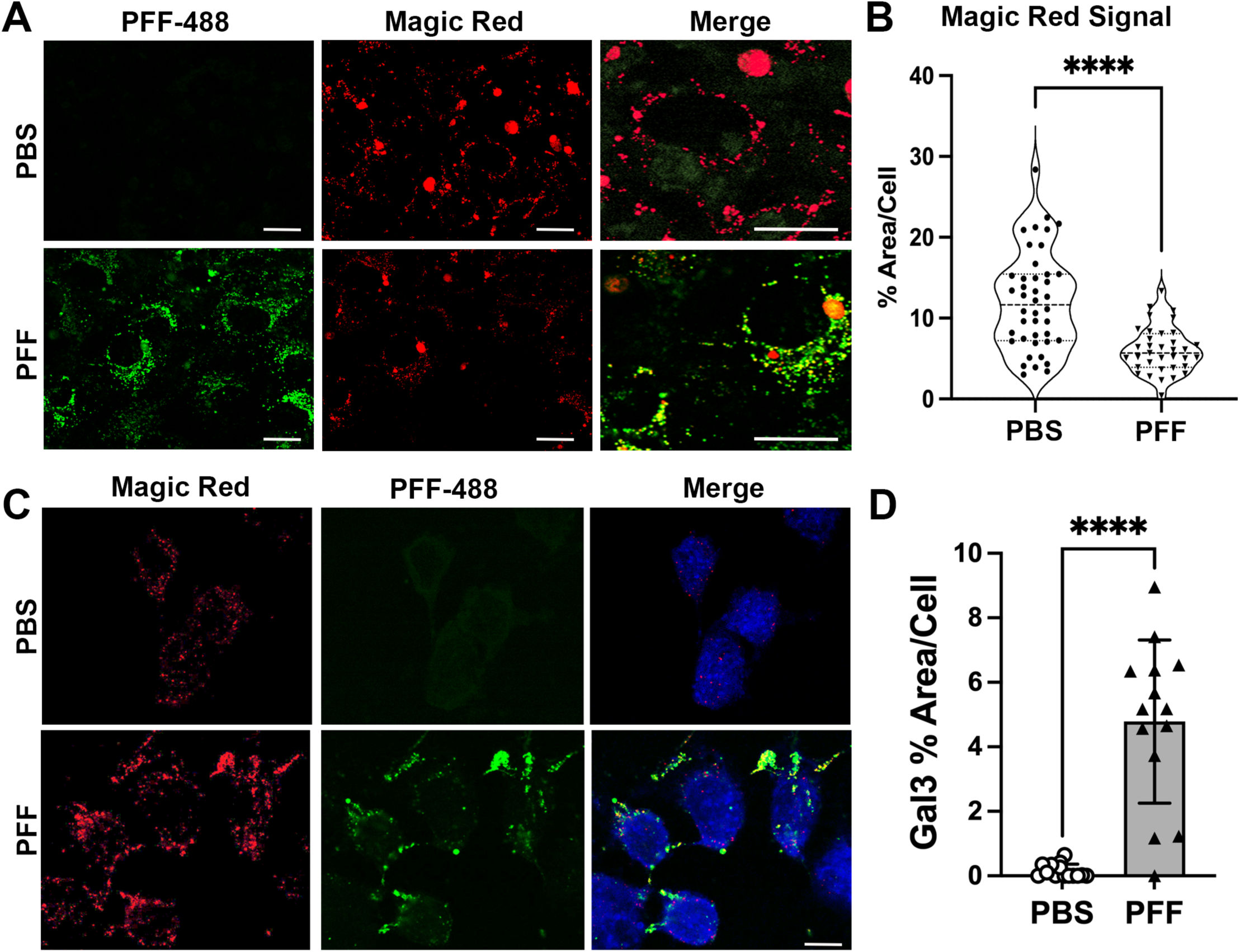
Internalized αS PFF inhibits lysosome function in neuronal cells. **A, B)** Neuronally differentiated CLU198 cells were treated with PFF-AF488, washed, and incubated for 24 h prior to confocal live cell imaging. The cells were also treated with Magic Red Cathepsin-B assay reagent for the last hour prior to imaging. (**A**) Representative confocal live cell images of PFF-AF488 (Green) and Magic Red (Red). Merge shows higher magnification to show details. (**B**) Violin Plot of % area/cell covered by Magic Red signal. The overall Magic Red signal, representing Cathepsin B activity, is significantly lower in PFF treated cells. **C, D**) Neuronally differentiated CLU198 cells were treated with PFF-AF488, washed, and incubated for 24-h, fixed in 4% PFA, and stained for Gal3, an indicator of lysosomal damage. The cells were imaged using confocal microscopy. Representative confocal images show increased Gal3 staining in PFF-treated cells (**C**). Gal3 percentage (%) of area per cell is significantly higher in PFF treated cells (**D**). \*\*\*\**p<0.0001*, unpaired *t*-test. Bar=10 μm.

We also used Galectin-3 (Gal3) staining as a marker of lysosomal integrity [31]. Gal3 is normally localized diffusely in the cytosol but when the lysosome becomes permeable/damaged, Gal3 translocates into the lysosome and exhibits punctate staining. Analysis of cells treated with PBS or αS PFF shows that αS PFF treatments lead to a significant increase in Gal3 staining (**Fig. 7C, D**). Finally, we analyzed lysosomal function in PCN using DQ-Red-BSA, which is targeted to lysosomes and resulting lysosomal proteolysis leads to an intense florescence signal [4]. In PCN, PFF treatment leads to a significant decrease in the DQ-Red fluorescence (**Fig. S7D, F**).

We also evaluated the involvement of other possible proteolytic machinery in the metabolism of αS PFF. Given the important relationship between the lysosomes and autophagy, we examined the possible role of autophagy in the metabolism of αS PFF. We examined the role of autophagy on αS PFF metabolism by inhibiting autophagy via the 3-methyladenine (3MA) treatment [32] (**Fig. 8A-C; Fig. S8A-D**) and promoting autophagy via rapamycin treatment [33] (**Fig. 8D-F; Fig. S8E, F**). 3MA treatment of differentiated CLU198 cells led to modest increases in αS levels at 3 h and 12 h following αS PFF treatment but the increase was not statistically significant (**Fig. 8A-C**). Non-neuronal cells treated with 3MA showed no effect on αS levels in astrocytes (**Fig. S8A, B**) and a modest increase in αS levels in HEK293 cells at 12 h following PFF treatment (**Fig. 8C, D**). Stimulation of autophagy by rapamycin treatment did not lead to any obvious alternations in the levels of αS following PFF treatment of differentiated CLU198 cells (**Fig. 8D-F**) and non-neuronal HEK293 cells (**Fig. S8F**). Despite the lack of rapamycin on αS PFF metabolism, we observe that rapamycin treatment clearly inhibited mTOR (pS6, 4EBP1) and increased autophagy (LC3 and p62) (Fig. S8E, F).

**Figure 8.**
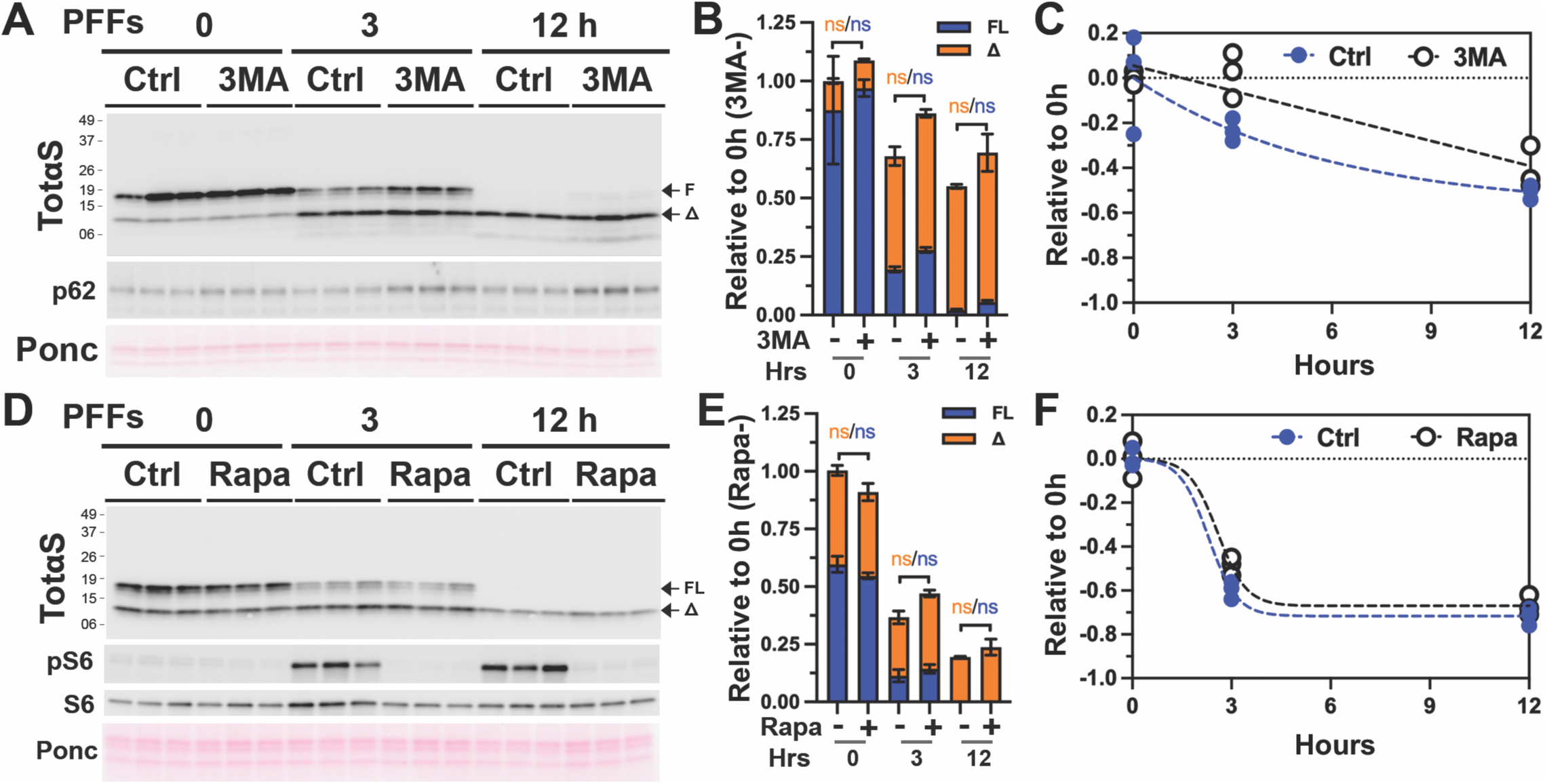
Autophagy is not a major factor in the metabolism of internalized αS PFF in neuronal cells. Differentiated hippocampal CLU198 cells and were pre-treated for 4h with an autophagy inhibitor 3 methyladenine (100 mM, 3MA, **A-C**) or an autophagy activator rapamycin (100 ηM, Rapa, **D-F**). The cells were treated with 4 µg/ml αS PFF for the last 2 h of 3MA treatment and washed. Washed cells were provided with fresh media containing PBS (Control, Ctrl), 3MA or Rapa and then incubated for 0-, 3-, and 12-h before harvesting. Levels of TotαS were detected by immunoblot analysis (**A, D**). Ponceau S (Ponc) staining was used to verify equal protein loading. The bar (**B, E**) and line (**C, F**) graphs show the relative levels of full-length (FL) and truncated (Δ) αS levels over time. Mean±SEM; n=3. n.s., not significant.

Analysis of the levels of active lysosomal enzyme cathepsin D shows that 3MA treatment decreases active cathepsin D levels in astrocytes and HEK293 cells (**Fig. S8B, D**). Thus, we believe 3MA treatment could be inhibiting the metabolism of αS PFF partly via induction of modest lysosomal deficit. Regardless, 3MA impacted αS PFF metabolism less than with the direct inhibition of lysosomes. Collectively, our results suggest that autophagy is a minor component in the regulation of αS PFF metabolism in neuronal cells. We also examined the potential role of the proteasome in αS PFF degradation in CLU198 and HEK-293 cells using, PS-341, a selective proteasomal inhibitor [34] (**Fig. S8 G, H)**. We found that proteasome inhibition did not affect the metabolism of internalized αS PFF (**Fig. S8 G, H**). In conclusion, our studies indicate that lysosome is the major degradation machinery for the internalized αS PFFs.

## Discussion

Progression of αS pathology in the brain is thought to involve cell-to-cell spreading of αS pathology where the pathogenic αS from doner neurons induces αS pathology in the neighboring neurons that internalize the pathogenic αS. In addition to neurons, glial cells can efficiently internalize extracellular αS variants released by neurons and may impact the spread of αS pathology. For example, astrocytes or microglia can attenuate the development of αS pathology in neurons by competing for the uptake of pathogenic αS [6,12,13,35]. However, most of the current studies examined the fate of exogenous αS at several hours or days following initial uptake. Thus, information about the short-term metabolism of the αS in various brain cell types is incomplete. To gain further insights about the roles of different brain cell types in the development of αS pathology, we examined how αS aggregates, in the form of αS PFF, are metabolized by various neural cells within hours of uptake rather than days. Our results show that neurons are inefficient in degrading internalized αS PFF and αS PFF stably accumulates as C-terminally truncated species. In neurons, we show that both the degradation and truncation of internalized αS PFF occurs in the endosome/lysosome compartment within minutes and hours following initial internalization of αS PFF. Moreover, truncation and stable accumulation of internalized αS PFF occurs in both primary cultured neurons as well as neuronal cell lines. Significantly, in neuronal cell lines (CLU-198 and SH-SY5Y), stable accumulation of truncated αS PFF is selectively associated with neuronal differentiation as undifferentiated cells rapidly degrade internalized αS PFF. Similarly, astrocytes and microglia rapidly degrade internalized αS PFF within 6-8 hours while αS PFF accumulates as truncated species in oligodendrocytes. Epitope mapping of the internalized αS PFF shows that the major 11.5 kDa truncated species, αS^Δ^, retains the N-terminal epitopes but presumably missing ∼25 amino acids from the C-terminal region. We also show that the lysosome is a major organelle responsible for the degradation of internalized αS PFF as inhibition of lysosomes significantly increased accumulation of αS PFF in all cell types. Finally, we show that in vivo, exogenous αS fibrils injected into the hippocampus are rapidly taken up and cleared by the glial cells while neurons exhibit slower but longer-term accumulation of αS fibrils.

We originally showed that a significant fraction of αS is normally truncated at the C-terminal region and the C-terminally truncated αS can promote αS aggregation [14]. Moreover, the abundance of C-terminally truncated αS is increased in PD cases as well as in the αS aggregates in vivo [14]. However, the current report indicates that the truncation of internalized αS PFF is different than the truncation of αS expressed in cells or αS aggregates extracted from human and mouse cells/brain. While ∼5-25% of endogenously expressed αS monomers accumulate as C-terminal truncated forms [14], internalized αS monomer is efficiently degraded without any accumulation of truncated αS. Further, αS aggregates extracted from mouse or human brains are partially truncated (∼25-50%) [14], while virtually 100% of internalized αS PFF are truncated by neurons. Similarly, analysis of lysosomes isolated from the brains of a TgA53T mouse line shows that αS^FL^ is more abundant than the truncated αS [28]. Direct comparison of truncated variants in αS PFF treated neurons with the αS aggregates extracted from the human PD case and the TgA53T model show that, with the *in vivo* derived αS aggregates, the major C-terminally truncated species resolve at ∼12 kDa [14], rather than ∼11.5 kDa seen with αS PFF **(Fig. 4G).** While we have not defined actual site of αS truncation, other studies have shown that several lysosomal proteases can truncate αS at several C-terminal residues [28], including Glu114 and Asn103 [26,28,36]. Interestingly, cleavage at Glu114 is resistant to Cathepsin inhibition [36], and cleavage at Asn103, while catalyzed by asparagine endopeptidase (AEP), is independent of lysosomal pH [37]. Based on the size of the αS^Δ^, it is unlikely that αS^Δ^ is truncated at Asn103. While the report by Quintin and colleagues [36] did not examine differential metabolism of αS fibrils in different cells types, we believe it is likely that αS^Δ^ observed in neurons is same as the αS truncated at Glu114 in HEK293 cells [36]. We do observe αS^Δ^ in HEK cells treated with both αS PFF and lysosomal inhibitor (**Fig. S6C**). Thus, we believe it is likely that very high levels of αS PFF (∼14 ug/ml) used in the Quintin et al study could be associated with lysosomal deficits. Overall, we provide a new observation that while astrocytes and microglia can rapidly degrade αS PFF without accumulation of αS^Δ^, oligodendrocytes resemble neurons as both cell types accumulate truncated αS^Δ^. Thus, C-terminal truncation and accumulation of exogenous αS PFF is a common feature of the cell types that are known to develop αS aggregates in human α-synucleinopathies.

A variety of other studies have shown that internalized αS or αS fibrils are targeted to lysosomes [4,23–25,38,39] and causes lysosomal deficits [4,9,23,40,41]. We have extended these studies by showing that internalized αS fibrils are differentially metabolized in neurons compared to non-neuronal cells. In neurons, we show that internalized αS fibrils are C-terminally truncated in endo-lysosomal compartment and stably accumulates. In contrast, glial cells rapidly degrade internalized αS fibrils. Further, we observe that similar kinetics of αS metabolism can be seen in vivo. Finally, we note that many of the prior studies on αS fibril metabolism utilize continuous treatment of the cells with very high levels of αS fibrils (0.7-1 μM, 8μg/ml-15μg/ml), which can artificially overwhelm the endo-lysosomal compartment, while we utilize much lower levels of αS fibrils (≤0.14 μM, ≤2 μg/ml).

Significantly, internalized αS PFF, even when extensively truncated, remains insoluble for at least 7 days, indicating that αS PFF remains aggregated in neurons for an extended period. The increased stability of the αS PFF in neurons may be because internalized αS PFF causes lysosomal dysfunction. However, this effect must not be a global lysosomal deficit but at an individual lysosome level since even very low levels of αS PFF are truncated and remain stable for an extended period. This result also indicates that even with the internalization of small amounts αS PFF by neurons, there is an extended timeframe for the internalized αS PFF to seed αS aggregation.

Collectively, we show that under normal conditions, any exogenous αS monomers are rapidly metabolized by all brain cell types but αS PFF, representing αS aggregates, are rapidly metabolized by glial cells but not by neurons. Our results suggest that studies on the cellular effects of αS PFF will need to consider the cell types used. We also predict that, if αS oligomers/aggregates are released extracellularly, astrocytes and microglia will efficiently remove the αS oligomers/aggregates under normal conditions, preventing significant transmission of αS oligomers/aggregates to neighboring neurons. However, under conditions that may lead to reduced αS uptake or lysosomal dysfunction, such as aging or increased inflammation, reduced metabolism of exogenous αS by glial cells likely promotes neuronal uptake of αS oligomers/aggregates and subsequent development of αS pathology.

## Materials and Methods

### Primary cell culture

Mouse pups between postnatal day 0-2 (P0-2) were used to establish primary cultures of cortical neurons and glia were established as previously described [42,43] (For protocol see: dx.doi.org/10.17504/protocols.io.14egn7jdmv5d/v1). For *Primary Cortical Neuron (PCN)*, dissociated cells from newborn mouse cortex were plated onto a Matrigel-coated cell culture dish using plating medium (PM; DMEM, 1 mM Sodium pyruvate, Glutamax, Penicillin-streptomycin and FBS) and on the following day, replaced with NbActiv4 (BrainBits LCC) containing FdU mitotic inhibitor (8 µM final) to halt the growth of non-neuronal cells. Culture media was changed periodically twice a week until the cells became mature neurons [∼12 days in vitro (DIV)], and then started various treatments as indicated. For *Mixed Glia culture*, 12,000,000 dissociated cells from the newborn brain were plated on a Matrigel-coated T75 flask using PM. Media was changed twice in a week until the cells became confluent (∼1 week in culture) (For protocol on glial isolation see: dx.doi.org/10.17504/protocols.io.36wgqn6k3gk5/v1). *Primary Microglial cultures* were established from the confluent mixed glial culture in a T75 flask via shaking at 220 rpm, 37°C for 1 h. Floating microglia were pelleted by centrifugation (300×g for 5 min). The cells were resuspended in PM and filtered through a 70 µm cell strainer. Cells were plated for 48 hours before the experiment. *Primary oligodendrocyte cultures* were established from the mixed glial culture depleted of microglia. Following microglial depletion, media was replaced and placed on a shaker (220 rpm, 37°C, overnight). Floating oligodendrocytes were filtered using a 40 µm cell strainer and spun down at 200×g for 10 min. Cells were resuspended in OPC media (PM + 50 µg/ml apo-transferrin, 5 µg/ml insulin, 30 ηM sodium selenite, 10 ηM D-biotin, 10 ηM hydrocortisone, 20 ηg/ml PDGF-AA and 20 ηg/ml bFGF). Cells were plated for 7-10 days before the experiment. *Primary Astrocyte cultures* were established following the removal of microglia and oligodendrocytes from the T75 flask. The remaining attached cells, representing astrocytes, were washed twice with PBS and detached using 0.25% Trypsin-EDTA, 5 ml NbAstro media was added and filtered through a 40 µm cells strainer and spun down at 300×g for 10 min. The pellet was resuspended in NbAstro media and filtered through 70 and then 40 µm cells strainer. Counted and cells were plated at 800,000 or 400,000 cells per well for 2-4 days before the experiment for biochemical or immunocytochemical analysis, respectively.

### Cell lines

SH-SY5Y human neuroblastoma (dx.doi.org/10.17504/protocols.io.rm7vzj6r2lx1/v1), CLU198 mouse hippocampal neuronal cell lines (dx.doi.org/10.17504/protocols.io.bp2l628wrgqe/v1), BV2 mouse microglia cell lines, and HEK-293 human embryonic kidney cell lines (dx.doi.org/10.17504/protocols.io.x54v92741l3e/v1) were used in this study. All these cells were growing on full medium (DMEM, 10% FBS and 1% Pen-Strep). For differentiation of SH-SY5Y full medium was exchanged with differentiation media containing Neurobasal-A, pen-strep, B27, and retinoic acid. And for CLU the differentiation media contained Neurobasal-A, pen-strep, B27, and glutamax. Cells were differentiated for at least one week before being used for experiments.

### Generation of recombinant α-synuclein (αS) pre-formed fibril (PFF)

Recombinant wild-type human/mouse α-Synuclein (αS) isolation, purification, and fibril formation were done as previously described [44] with modifications (For protocol see: dx.doi.org/10.17504/protocols.io.81wgbxjx3lpk/v1)[42]. Purified αS monomers were used to assemble αS PFF by agitation as previously described [42]. The PFF was diluted in PBS to 5 μg/μl aliquot and kept in a −80°C freezer. For internalization and processing study, PFF from frozen stock was diluted to 0.25 µg/µl in PBS, sonicated (1 Sec “on” then 1 Sec “off”) for 120

Sec with 20% amplitude by utilizing a Fisher Scientific Branson micro probe-tip sonicator (Fischer Scientific; Hampton, NH). Then, the sonicated PFFs were added into the media for the indicated time and doses (4 µg/ml, if otherwise indicated) to study primary cortical neurons, primary microglia, primary astrocytes, primary oligodendrocytes, hippocampal cell line CLU, SHSY5Y and HEK cells.

### α-synuclein (αS) preformed fibrils (PFF) uptake and degradation/clearance assays

For the uptake assay (For protocol see: dx.doi.org/10.17504/protocols.io.e6nvw1e79lmk/v1), cells were treated with fresh cultured media containing 4 µg/ml of αS PFF and incubated for the indicated duration. Total lysates of fibril-treated cells were harvested at the end of the indicated time points.

For the clearance assay (dx.doi.org/10.17504/protocols.io.n2bvjndmxgk5/v1), cells were pretreated with 4 µg/ml of αS PFF for 2 h, then washed with PBS or PBS supplemented with trypsin (0.005%) for 1 min to remove any excess PFFs bound on the external cell surface. The washed cells were incubated with fresh media in the presence or absence of other drugs/inhibitors treatment. Cells were harvested at the indicated time points.

### Subcellular lysosomal fractionation

After the designated treatment, cells were washed thrice with DMEM and extracted in 0.5 ml of homogenization buffer (250 mM sucrose, 2 mM EDTA, 1.5 mM magnesium chloride, 10 mM potassium chloride, and 20 mM HEPES) supplemented with proteinase and phosphatase inhibitors (For protocol see: dx.doi.org/10.17504/protocols.io.kqdg32kdqv25/v1). The cells were gently detached using a cell scraper and homogenized using a Teflon homogenizer (12 strokes). 50 µl of total homogenates (TH) were separated and lysed with TNE lysis buffer. The rest of the TH was centrifuged at 1000×g for 10 minutes at 4°C. The resulting supernatant was centrifuged for 20,000×g for 20 minutes at 4°C to collect the precipitate as a crude lysosomal fraction (CLF). The CLF was lysed with TNE lysis buffer and used for Western blot analysis.

### Triton X-100 fractionation of soluble and insoluble αS

Protein extraction and fractionation into total lysates (TL), Triton X-100 soluble (S), and Triton X-100 Insoluble (IS) fraction was conducted as described previously (For protocol see: dx.doi.org/10.17504/protocols.io.5qpvob21zl4o/v1)[14,42]. Briefly, cells were washed using cold PBS, TNE+1% TX-100 was added on ice for 5 mins and sonicated to achieve the TL fractions. The TL was centrifuged on the Airfuge (Beckman Coulter) at 100,000xg for 10 minutes. The supernatant was adjusted to complete TNE lysis buffer and considered as a soluble fraction. Washed pellets were re-suspended in complete TNE lysis buffer as the insoluble fractions.

### Protein extraction and Immunoblot analysis of protein expression

Cells were lysed in TNE lysis buffer (50 mM Tris, 150 mM NaCl, 5 mM EDTA adjusted to pH at 7.4) containing 1% SDS, 0.5% NP-40, 0.5% DOC and protease/phosphatase inhibitors as previously described (For protocol see: dx.doi.org/10.17504/protocols.io.3byl4qpqjvo5/v1)[10]. Immunoblot analysis was conducted as described previously [10,42,45]. Briefly, relative protein levels of Total αS (SYN-1), Hu Specific αS, NAC-2 αS, N-2 αS, p62, LC3 and other proteins were determined from cell extracts by quantitative immunoblots analysis using chemiluminescence detection of horseradish peroxidase-conjugated secondary antibodies on the GE Image-Quant LAS-4010 (GE Healthcare, Waukesha, WI). Image-Quant software (GE; RRID:SCR_014246; https://www.cytivalifesciences.com) was used to determine the intensity of the immunoreactive bands [10,42,45].

### Labeling of α-synuclein pre-formed fibril with Alex-Fluor 488 (PFF-AF-488)

Alex Fluor 488 Microscale Protein Labelling Kit (AF-488, A30006) (Invitrogen) was used to label PFFs according to the manufacturer’s protocol (dx.doi.org/10.17504/protocols.io.4r3l2q6rql1y/v1). AF-488 reactive dye has a tetrafluorophenyl (TFP) ester moiety that is more stable in solution than the commonly used succinimidyl (NHS) ester. TFP ester reacts efficiently with primary amines of protein to form a stable dye-protein conjugate and is independent of pH between 4 and 10. Hu-WT-αS fibrils (PFFs) were diluted at a concentration of 1µg/µL in PBS followed by sonication (1 Sec on then 1 Sec off) for 120 Sec with 20% amplitude. Sonicated PFFs were used to label with AF-488.

### Immunocytochemistry

Immunocytochemistry was conducted as described previously [10,42,43] (For protocol see: dx.doi.org/10.17504/protocols.io.36wgqnorygk5/v1). Briefly, cells grown on coverslips were fixed in 4% paraformaldehyde (PFA) and immunostained followed by confocal imaging. Alex fluor (AF-647 or AF-488) conjugated secondary antibody was used.

### Analysis of αS PFF-AF-488 uptake in vivo following intrahippocampal Injections of αS PFF-AF-488

AF-488-conjugated αS PFF described above were injected into to covalently label 5-month-old α-synuclein knockout mice (*snca*^-/-^; n=3 per timepoint). The mice were anesthetized using 3% isoflurane mixed with oxygen, placed on a stereotaxic frame, and the mice received a unilateral injection of 2.5 μL total volume per depth (185 ng/µl)) using a 10 μL syringe (Hamilton; Reno, NV, USA) at a rate of 200 nL/min into the hippocampus and overlying cortex (coordinates: +2.5 mm mediolateral; −2.4 mm anteroposterior; −2.4 mm followed by −1.0 dorsoventral from the skull). Injections were followed by a 5 min resting period prior to removal of the needle. The brains were collected following 3 hours and 24 hours post injection.

For brain collection, the mice were anaesthetized with isoflurane and intracardially perfused with potassium-free phosphate-buffered saline (PBS) followed by 4% paraformaldehyde (PFA) in PBS. Brains were removed and postfixed in PFA for 24 h, cryoprected in 30% sucrose solution, and 20 µm thick free-floating coronal sections were obtained using a freezing sliding microtome. The sections were immunostained with various primary antibodies followed by confocal microscopy. PFF-488 and lysosomal marker cathepsin-D (Cath-D) were colocalized with various cellular markers: Astrocytes, GFAP; Microglia, Iba1; Neuron, NeuN. PFF-488 trafficking study was done by monitoring its localization into sub-cellular level (Neuron, Astrocytes and Microglia) as well as lysosomal organelles level of brain Hippocampus area.

In this study we focused on CA1, Molecular Layer (ML) of Dentate Gyrus (DG) of Hippocampus. To quantify the cell-type colocalization, 40× images were used from multiple sections from each animal. The total number of cells and co-localized puncta were counted using HALO Quantitative Image Analyzer. Finally, the fraction of specific cellular co-localization of PFF-488 was calculated by dividing the number of PFF puncta with the total number of the cells. Co-localization of cell-specific PFF-488 with Cath-D expressed as the Manders coefficient estimated by Image-J software with JACop plugin.

### Analysis of Lysosome Function

To measure lysosomal Cathepsin B activity in live cells, we used the Magic Red Fluorescent Cathepsin B assay kit (Immunochemistry Technologies, CA; dx.doi.org/10.17504/protocols.io.ewov192b2lr2/v1). Briefly, cells were plated on coverslips and following PFF-488 uptake for indicated times, the cells were washed [PBS/trypsin (0.01%)] and treated with 1x Magic Red solution for 15 minutes at 37°C per manufacturer instruction. Live cell confocal imaging was done immediately by confocal microscopy where the lysosomal Cathepsin B activity is indicated by red fluorescence puncta produced by the hydrolysis of fluorogenic substrate, Magic Red. Images were taken from each culture using identical conditions and the confocal images were used to analyze the Magic Red signal in each cell using Image J. Briefly, each cell was outlined and the % area of the cell covered by Magic Red signal, as well as total signal intensity per cell, was determined.

To measure lysosomal membrane permeabilization (LMP), we immunostained for Galactin-3 (Gal3) [31] (dx.doi.org/10.17504/protocols.io.eq2lywxbevx9/v1). Gal3 is a sugar-binding protein that translocates from cytosol to lysosome when the membrane becomes permeable due to lysosomal damage. Once LMP happens, the Gal3 fluorescence pattern changes from a diffuse condition to a dotted punctate structure. For the LMP assay, cells were plated on coverslips and treated with PFF-488 as indicated time. Cells were fixed in 4% PFA and immunostained using Gal3 primary and AF-647 conjugated secondary antibodies. Images were taken by confocal microscopy. To measure Gal3 accumulation in permeable lysosomes, punctate Gal3 staining was determined using Image J. Briefly, each cell was outlined and the % area of the cell covered by Gal3 immunoreactivity, as well as total signal intensity per cell, was determined.

To measure overall lysosomal function in PCN, we used the DQ-Red-BSA assay (dx.doi.org/10.17504/protocols.io.j8nlk8dp6l5r/v1). Briefly, cells were cultured and treated with PFF-488 as mentioned above. Following indicated treatments, 10 µg/ml of DQ-BSA Red (Thermo Fisher Scientific) was dissolved in cell culture media and incubated for the last 90 minutes at 37°C. Fixed cells using 4% PFA at room temperature and images were taken following identical conditions throughout the treatment condition. DQ-BSA Red fluorescent signal is detected only when the fluorogenic substrate is hydrolyzed by lysosomal proteases. The red fluorescence resulting from lysosomal targeting of DQ-Red-BSA was determined using Image J. Because of an extensive network of neurites with DQ-Red signal in neurons, we quantified total DQ-Red signal (% area covered and total signal intensity) per microscopic field in at least 4 independent areas. The total % area covered by the DQ-Red signal was divided by the number of cells (DAPI stained nuclei) in the field analyzed.

### Antibodies

The primary antibodies used in this study is listed in the supplementary table (**Table S1)**.

### Statistical analysis

To test for statistical significance between treatment groups, data was analyzed by one-way or two-way analysis of variance (ANOVA) followed by a Multiple Comparison post hoc test (Tukey’s/Dunnett’s/Bonferroni’s), or Student’s t-test. All tests were performed using GraphPad PRISM Software (Version 10; RRID:SCR_002798; https://www.graphpad.com). All the data are expressed as means ± S.E. Probability (p) values less than 0.05 were considered significantly different.

## Ethics Statement for Animal Research

All animal studies were performed in accordance with the national ethics guidelines for the use of animals in research and approved by the Institutional Animal Care and Use Committee (IACUC) at the University of Minnesota.

## Acknowledgments

We thank Drs. Mohammad Abdur Rashid and Balvindar Singh for their early preliminary observations leading to the current report. We also thank Mr. Hector Martell Martinez for help with glial cultures.

## Availability of data and materials

The datasets used and/or analyzed as well as a table of key resources (KRT) for the current study are available on Zenodo (<u>10.5281/zenodo.12522122</u>). Any additional data and materials are available from the corresponding author upon reasonable request.

## Author Contributions

M.R.K. and M.K.L. conceived and designed the study. All authors were responsible for the acquisition of the data. M.R.K., S.C.V., and M.K.L. were responsible for data analysis and writing/editing of the manuscript.

## Abbreviations

Baf A1: bafilomycin A1
DMEM: Dulbecco’s modified Eagle’s medium
DMSO: Dimethyl sulfoxide
GAPDH: glyceraldehyde 3-phosphate dehydrogenase
LC3: microtubule-associated protein 1A/1B-light chain 3
NAC domain: non-amyloidal component
PD: Parkinson’s diseases
p62 (SQSTM1): sequestosome 1
PFF: αS pre-formed-fibril
PCN: primary cortical neuron
PMG: primary microglia
αS: α-synuclein
αS^ΔC^: C-terminally truncated αS
FL: full-length

## References

[1] Spillantini MG, Schmidt ML, Lee VM, et al. Alpha-synuclein in Lewy bodies. Nature. 1997 Aug 28;388(6645):839–40.

[2] Cremades N, Cohen SI, Deas E, et al. Direct observation of the interconversion of normal and toxic forms of alpha-synuclein. Cell. 2012 May 25;149(5):1048–59.

[3] Bengoa-Vergniory N, Roberts RF, Wade-Martins R, et al. Alpha-synuclein oligomers: a new hope. Acta Neuropathol. 2017 Dec;134(6):819–838.

[4] Senol AD, Samarani M, Syan S, et al. alpha-Synuclein fibrils subvert lysosome structure and function for the propagation of protein misfolding between cells through tunneling nanotubes. Plos Biol. 2021 Jul;19(7).

[5] Bae EJ, Yang NY, Song M, et al. Glucocerebrosidase depletion enhances cell-to-cell transmission of alpha-synuclein. Nat Commun. 2014 Aug 26;5:4755.

[6] Loria F, Vargas JY, Bousset L, et al. alpha-Synuclein transfer between neurons and astrocytes indicates that astrocytes play a role in degradation rather than in spreading. Acta Neuropathol. 2017 Nov;134(5):789–808.

[7] Chung HK, Ho HA, Perez-Acuna D, et al. Modeling alpha-Synuclein Propagation with Preformed Fibril Injections. J Mov Disord. 2019 Sep;12(3):139–151.

[8] Volpicelli-Daley LA, Luk KC, Patel TP, et al. Exogenous alpha-synuclein fibrils induce Lewy body pathology leading to synaptic dysfunction and neuron death. Neuron. 2011 Oct 6;72(1):57–71.

[9] Hoffmann AC, Minakaki G, Menges S, et al. Extracellular aggregated alpha synuclein primarily triggers lysosomal dysfunction in neural cells prevented by trehalose. Scientific reports. 2019 Jan 24;9(1):544.

[10] Karim MR, Liao EE, Kim J, et al. alpha-Synucleinopathy associated c-Abl activation causes p53-dependent autophagy impairment. Mol Neurodegener. 2020 Apr 16;15(1):27.

[11] Poehler AM, Xiang W, Spitzer P, et al. Autophagy modulates SNCA/alpha-synuclein release, thereby generating a hostile microenvironment. Autophagy. 2014;10(12):2171–92.

[12] Scheiblich H, Dansokho C, Mercan D, et al. Microglia jointly degrade fibrillar alpha-synuclein cargo by distribution through tunneling nanotubes. Cell. 2021 Sep 30;184(20):5089–5106 e21.

[13] Choi I, Zhang Y, Seegobin SP, et al. Microglia clear neuron-released alpha-synuclein via selective autophagy and prevent neurodegeneration. Nat Commun. 2020 Mar 13;11(1):1386.

[14] Li W, West N, Colla E, et al. Aggregation promoting C-terminal truncation of alpha-synuclein is a normal cellular process and is enhanced by the familial Parkinson’s disease-linked mutations. Proc Natl Acad Sci U S A. 2005 Feb 8;102(6):2162–7.

[15] Li W, Lesuisse C, Xu Y, et al. Stabilization of alpha-synuclein protein with aging and familial parkinson’s disease-linked A53T mutation. J Neurosci. 2004 Aug 18;24(33):7400–9.

[16] Gingerich S, Kim GL, Chalmers JA, et al. Estrogen receptor alpha and G-protein coupled receptor 30 mediate the neuroprotective effects of 17beta-estradiol in novel murine hippocampal cell models. Neuroscience. 2010 Sep 29;170(1):54–66.

[17] Mahul-Mellier AL, Burtscher J, Maharjan N, et al. The process of Lewy body formation, rather than simply alpha-synuclein fibrillization, is one of the major drivers of neurodegeneration. Proc Natl Acad Sci U S A. 2020 Mar 3;117(9):4971–4982.

[18] Peng C, Gathagan RJ, Covell DJ, et al. Cellular milieu imparts distinct pathological alpha-synuclein strains in alpha-synucleinopathies. Nature. 2018 May;557(7706):558–563.

[19] Reddy K, Dieriks BV. Multiple system atrophy: alpha-Synuclein strains at the neuron-oligodendrocyte crossroad. Mol Neurodegener. 2022 Nov 26;17(1):77.

[20] Halliday GM, Holton JL, Revesz T, et al. Neuropathology underlying clinical variability in patients with synucleinopathies. Acta Neuropathol. 2011 Aug;122(2):187–204.

[21] Luth ES, Stavrovskaya IG, Bartels T, et al. Soluble, prefibrillar alpha-synuclein oligomers promote complex I-dependent, Ca2+-induced mitochondrial dysfunction. J Biol Chem. 2014 Aug 1;289(31):21490–507.

[22] Perrin RJ, Payton JE, Barnett DH, et al. Epitope mapping and specificity of the anti-alpha-synuclein monoclonal antibody Syn-1 in mouse brain and cultured cell lines. Neurosci Lett. 2003 Oct 2;349(2):133–5.

[23] Masaracchia C, Hnida M, Gerhardt E, et al. Membrane binding, internalization, and sorting of alpha-synuclein in the cell. Acta Neuropathol Commun. 2018 Aug 14;6(1):79.

[24] Rodriguez L, Marano MM, Tandon A. Import and Export of Misfolded alpha-Synuclein. Front Neurosci. 2018;12:344.

[25] Burbidge K, Rademacher DJ, Mattick J, et al. LGALS3 (galectin 3) mediates an unconventional secretion of SNCA/alpha-synuclein in response to lysosomal membrane damage by the autophagic-lysosomal pathway in human midbrain dopamine neurons. Autophagy. 2022 May;18(5):1020–1048.

[26] Zhang Z, Kang SS, Liu X, et al. Asparagine endopeptidase cleaves alpha-synuclein and mediates pathologic activities in Parkinson’s disease. Nat Struct Mol Biol. 2017 Aug;24(8):632–642.

[27] Sorrentino ZA, Giasson BI. The emerging role of alpha-synuclein truncation in aggregation and disease. J Biol Chem. 2020 Jul 24;295(30):10224–10244.

[28] McGlinchey RP, Lacy SM, Huffer KE, et al. C-terminal alpha-synuclein truncations are linked to cysteine cathepsin activity in Parkinson’s disease. J Biol Chem. 2019 Jun 21;294(25):9973–9984.

[29] Lie PPY, Nixon RA. Lysosome trafficking and signaling in health and neurodegenerative diseases. Neurobiol Dis. 2019 Feb;122:94–105.

[30] Song Q, Meng B, Xu H, et al. The emerging roles of vacuolar-type ATPase-dependent Lysosomal acidification in neurodegenerative diseases. Transl Neurodegener. 2020 May 11;9(1):17.

[31] Aits S. Methods to Detect Loss of Lysosomal Membrane Integrity. Methods Mol Biol. 2019;1880:315–329.

[32] Wu YT, Tan HL, Shui G, et al. Dual role of 3-methyladenine in modulation of autophagy via different temporal patterns of inhibition on class I and III phosphoinositide 3-kinase. J Biol Chem. 2010 Apr 2;285(14):10850–61.

[33] Sarkar S, Ravikumar B, Floto RA, et al. Rapamycin and mTOR-independent autophagy inducers ameliorate toxicity of polyglutamine-expanded huntingtin and related proteinopathies. Cell Death Differ. 2009 Jan;16(1):46–56.

[34] Bazzaro M, Lin Z, Santillan A, et al. Ubiquitin proteasome system stress underlies synergistic killing of ovarian cancer cells by bortezomib and a novel HDAC6 inhibitor. Clin Cancer Res. 2008 Nov 15;14(22):7340–7.

[35] Tsunemi T, Ishiguro Y, Yoroisaka A, et al. Astrocytes Protect Human Dopaminergic Neurons from alpha-Synuclein Accumulation and Propagation. J Neurosci. 2020 Nov 4;40(45):8618–8628.

[36] Quintin S, Lloyd GM, Paterno G, et al. Cellular processing of alpha-synuclein fibrils results in distinct physiological C-terminal truncations with a major cleavage site at residue Glu 114. J Biol Chem. 2023 Jul;299(7):104912.

[37] Kang SS, Ahn EH, Zhang Z, et al. alpha-Synuclein stimulation of monoamine oxidase-B and legumain protease mediates the pathology of Parkinson’s disease. EMBO J. 2018 Jun 15;37(12).

[38] Lee HJ, Suk JE, Bae EJ, et al. Assembly-dependent endocytosis and clearance of extracellular alpha-synuclein. Int J Biochem Cell Biol. 2008;40(9):1835–49.

[39] Desplats P, Lee HJ, Bae EJ, et al. Inclusion formation and neuronal cell death through neuron-to-neuron transmission of alpha-synuclein. Proc Natl Acad Sci U S A. 2009 Aug 4;106(31):13010–5.

[40] Flavin WP, Bousset L, Green ZC, et al. Endocytic vesicle rupture is a conserved mechanism of cellular invasion by amyloid proteins. Acta Neuropathol. 2017 Oct;134(4):629–653.

[41] Freeman D, Cedillos R, Choyke S, et al. Alpha-synuclein induces lysosomal rupture and cathepsin dependent reactive oxygen species following endocytosis. PLoS One. 2013;8(4):e62143.

[42] Vermilyea SC, Christensen A, Meints J, et al. Loss of tau expression attenuates neurodegeneration associated with alpha-synucleinopathy. Transl Neurodegener. 2022 Jul 1;11(1):34.

[43] Nanclares C, Poynter J, Martell-Martinez HA, et al. Dysregulation of astrocytic Ca(2+) signaling and gliotransmitter release in mouse models of alpha-synucleinopathies. Acta Neuropathol. 2023 May;145(5):597–610.

[44] Volpicelli-Daley LA, Luk KC, Lee VM. Addition of exogenous alpha-synuclein preformed fibrils to primary neuronal cultures to seed recruitment of endogenous alpha-synuclein to Lewy body and Lewy neurite-like aggregates [Research Support, N.I.H., Extramural]. Nat Protoc. 2014 Sep;9(9):2135–46.

[45] Singh B, Covelo A, Martell-Martinez H, et al. Tau is required for progressive synaptic and memory deficits in a transgenic mouse model of alpha-synucleinopathy. Acta Neuropathol. 2019 Jun 6.

