## Supplemental Table S1 and Figure S1-S9 for "Internalized α-synuclein fibrils become truncated and resist degradation in neurons while glial cells rapidly degrade α-synuclein fibrils"

### Supplementary Information

Table S1

Figures S1-S9

Internalized $\alpha$-synuclein fibrils are rapidly truncated and resists degradation in neurons while glial cells rapidly degrade $\alpha$-synuclein fibrils.

Md. Razaul Karim, Emilie Gasparini, Elizabeth Tiegs, Riley Schlichte, Scott C. Vermilyea, and Michael K. Lee

| **Loading Controls** | **Host** | **Source** | **Reference** | **RRID** | **Use** |
|---|---|---|---|---|---|
| GAPDH (D16H11) | Rbt | Cell Signaling | 5174 | AB_10622025 | WB |
| α-tubulin | Rbt | Abcam | 4074 | AB_2288001 | WB |
| **α-Synuclein Species** | **Host** | **Company** | **Reference** | **RRID** | **Use** |
| α-Synuclein (total, Syn1) | Rt | BD Transduction | 610787 | AB_398108 | WB |
| HuαS | Rbt | In House | [1] | N/A | WB,ICC |
| LB509 | Ms | Abcam | 27766 | AB_727020 | WB/ICC |
| NAC-2 αS/N-2 αS | Rbt | Pekka Jäkälä, Kupio University | [1] | N/A | WB |
| Pan-Syn | Rbt | Abcam | ab53726 | AB_882803 | WB |
| **Glial and Neuronal Markers** | **Host** | **Company** | **Reference** | **RRID** | **Use** |
| Iba1 | Rbt | Wako Chemical | 019-19741 | AB_839504 | ICC |
| GFAP | Rbt | Dako Cytomation | Z0334 | AB_10013382 | ICC |
| NeuN | Ms | Millipore | MAB377 | AB_2313673 | ICC |
| O4 | MS | R&D System | MAB1326 | AB_357617 | ICC |
| **Autophagy/Lysosome Function/Organelle** | **Host** | **Company** | **Reference** | **RRID** | **Use** |
| LC3 | Rbt | Cell Signaling | 2775 | AB_915950 | WB, ICC |
| p62 | Rbt | Cell Signaling | 5114 | AB_10624872 | WB, ICC |
| pS6 | Rbt | Cell Signaling | 2211 | AB_331679 | WB |
| S6 (total) | Ms | Cell Signaling | 2317 | AB_2238583 | WB |
| 4EBP | Rbt | Cell Signaling | 9644 | AB_2097841 | WB |
| CTSD | Ms | Abcam | ab75852 | AB_1523267 | WB |
| Gal3 | Rbt | Abcam | ab2785 | AB_2291667 | ICC |
| Poly-Ubq | Rbt | Dako | Z0458 | AB_2315524 | WB |
| Grp78 | Rbt | Novus | NB300-520 | AB_10000968 | ICC |
| Lamp1 | Rat | LS-Bio | LS-B4246 | AB_10718424 | WB, ICC |
| Lamp2 | Rbt | LS-Bio | LS-B581 | AB_909691 | ICC |
| EEA1 | Rbt | Gene Tex | GTX 109638 | AB_1950162 | WB, ICC |
| GM-130 | Rbt | Novus | NBP2-53420 | AB_2916095 | ICC |
| P62 | Ms | Abcam | ab56416 | AB_945626 | ICC |
| LC3 | Rbt | Cell Signaling | 3868 | AB_2137707 | ICC |

**Table S1.** List of primary antibodies utilized in experiments. Ms, Mouse; Rt, Rat; Rbt, Rabbit; WB, western blot; ICC, immunocytochemistry.

1. Li, W., et al., *Aggregation promoting C-terminal truncation of alpha-synuclein is a normal cellular process and is enhanced by the familial Parkinson's disease-linked mutations.* Proc Natl Acad Sci U S A, 2005. **102**(6): p. 2162-7.

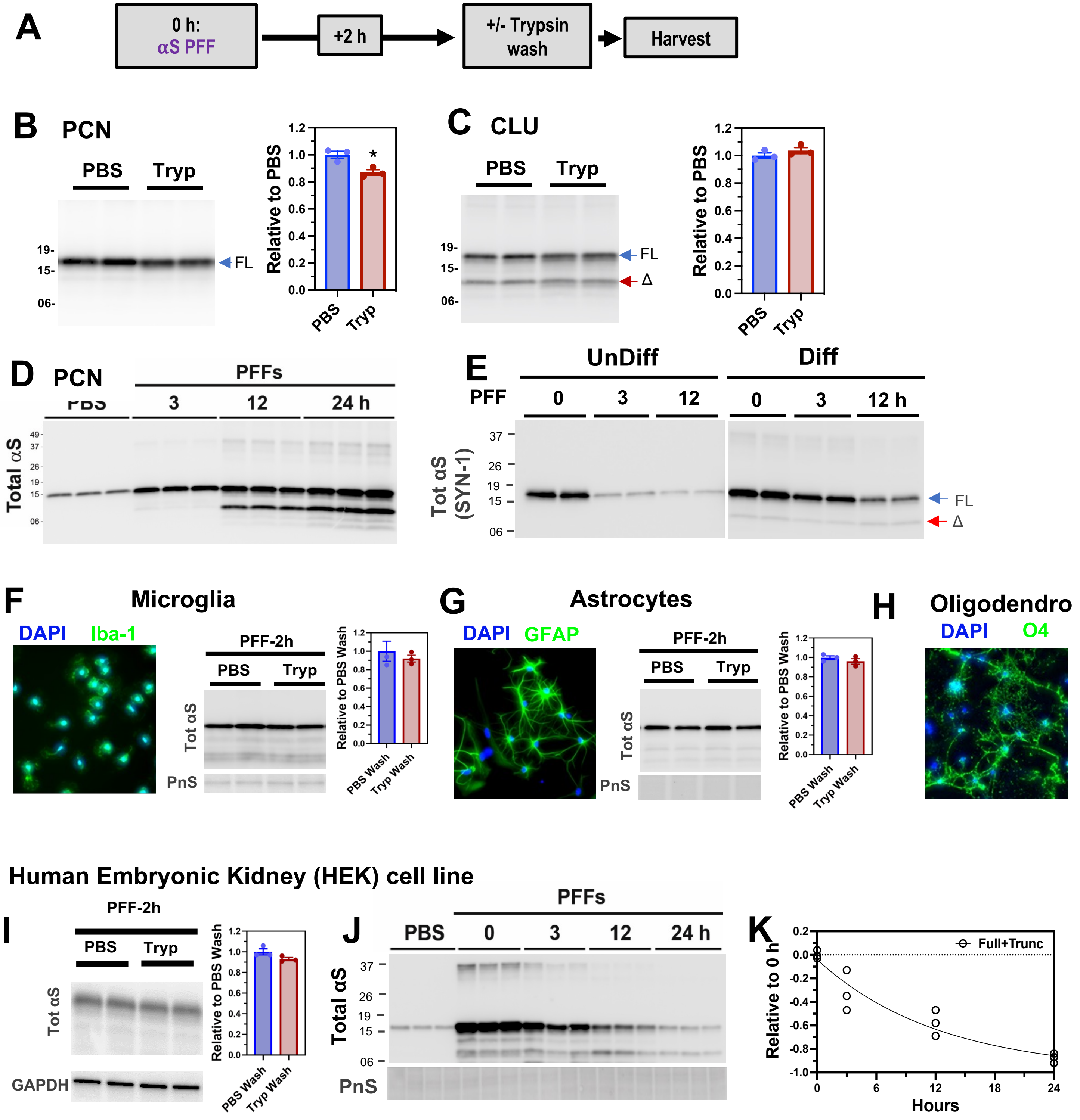


**Figure S1. A-E) Comparison of PBS wash and Trypsin wash in removing uninternalized αS PFF (A-D) and metabolism of αS PFF in SH-SY5Y cells (E). A)** Scheme showing cultured primary cortical neuron **(PCN, B)** or CLU198 cells **(CLU, C)** incubated with 4 μg/ml αS PFF for 2 hours and washed with PBS or trypsin before the harvest. The levels of Tot αS were detected by immunoblot analysis. **B)** In PCN, approximately ~85-90% residual αS following PBS wash is accounted by trypsin resistant fraction. **C)** In CLU cells, the amount of residual αS is not different between PBS and trypsin-washed cells. These results indicate that the bulk of residual αS following PBS wash is internalized αS (*$p<0.05$, t-test, Mean±SEM; n=3). **D)** Analysis αS PFF

**Figure S1. Continued**

uptake in PCN without washing shows that αS continues to increase past the 24 hours following PFF treatment. Thus, uptake of new αS PFF occurs for prolong periods without washing. **E)** Uptake and metabolism of αS PFF in an undifferentiated **(UnDiff)** and neuronally differentiated **(Diff)** human neuroblastoma cell line (SH-SY5Y cells). Immunoblot analysis of total lysates for αS shows that, similar to the CLU cells (Fig. 1G), internalized αS is rapidly degraded in undifferentiated SH-SY5Y cells. In differentiated SH-SH5Y cells, internalized αS is more stable with the presence of the truncated αS (Δ). **F-I) Uptake of αS pre-formed fibrils (PFFs) in glial cells and in HEK293 cells.** primary glial cells were isolated and cultured from newborn mouse pup's cortex and the purity was confirmed by staining with specific cellular marker Iba1 **(F)**, GFAP**(G)**, and O4 **(H)** antibodies for microglia, astrocytes, and oligodendrocytes, respectively. Micorglia **(F)**, Astrocytes **(G)**, and HEK293 **(I)** cells were also incubated with 4 μg/ml αS PFF for 2 hours followed by PBS or trypsin wash before harvesting. The levels of Tot αS were detected, (Mean±SEM; n=3). **J, K) Degradation of αS PFFs in HEK-293 cells.** Cells were pre-incubated for 2h with 4 μg/ml αS PFF followed by trypsin wash. After washing, cells were replenished with full media and incubated for the indicated time and levels of Tot αS were detected. The graph shows the amount of residual αS remaining from the 0 h time point. Ponceau S (PnS) protein stain or GAPDH were used to confirm loading.

Figure S2.

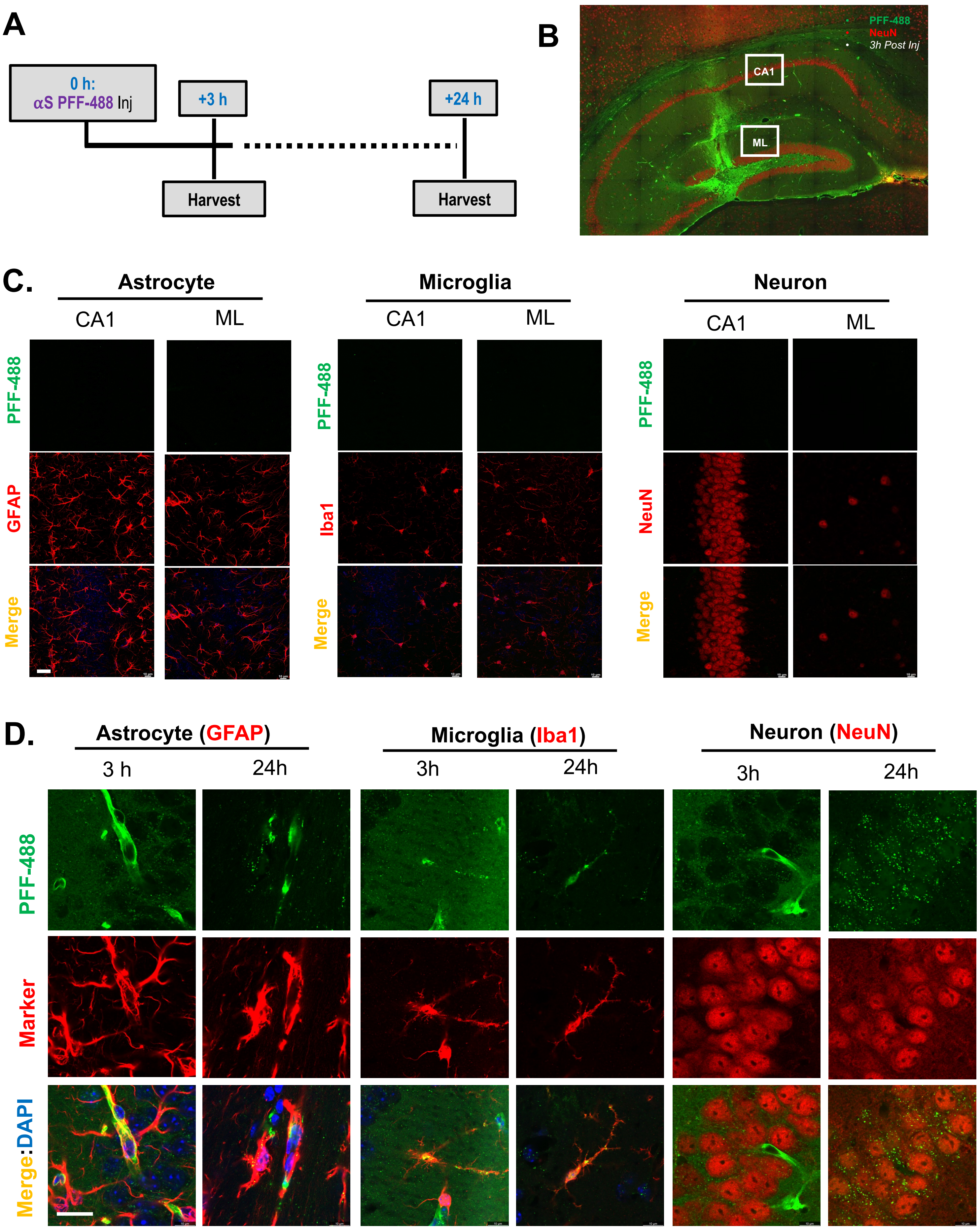

A
0 h:
αS PFF-488 Inj
+3 h
+24 h
Harvest
Harvest
B
PFF-488
NeuN
3h Post Inj
CA1
ML
C.
Astrocyte
Microglia
Neuron
CA1
ML
PFF-488
GFAP
Iba1
NeuN
Merge
D.
Astrocyte (GFAP)
Microglia (Iba1)
Neuron (NeuN)
3 h
24h
3h
PFF-488
Marker
Merge:DAPI

**Figure S2. Cellular localization of exogenous αS PFF injected into mouse brain. A)** Schematic for the analysis of AF488-labelled αS PFF injected into mouse brain. **B)** Low magnification immunofluorescence microscopic image showing αS PFF-488 (green) and NeuN (red) immunostaining of the brain sections from 3 hours post injection. Note the highly levels of αS PFF-488 staining at the injection site and along the needle tract. White rectangles shows the regions examined and quantified. CA1-hippocampal pyramidal cell layer, ML-molecular layer or dentate gyrus. **C)** Imaging of contralateral side for αS PFF-488 along with cell type markers (GFAP, Iba1, NeuN) at 3 hours post injection. No αS PFF-488 associated signal is seen. **D)** Confocal slice image showing that the αS PFF-488 and the cell type markers are colocalizaing with the same 1 μm plane. Bar=20μm.

### Figure S3.

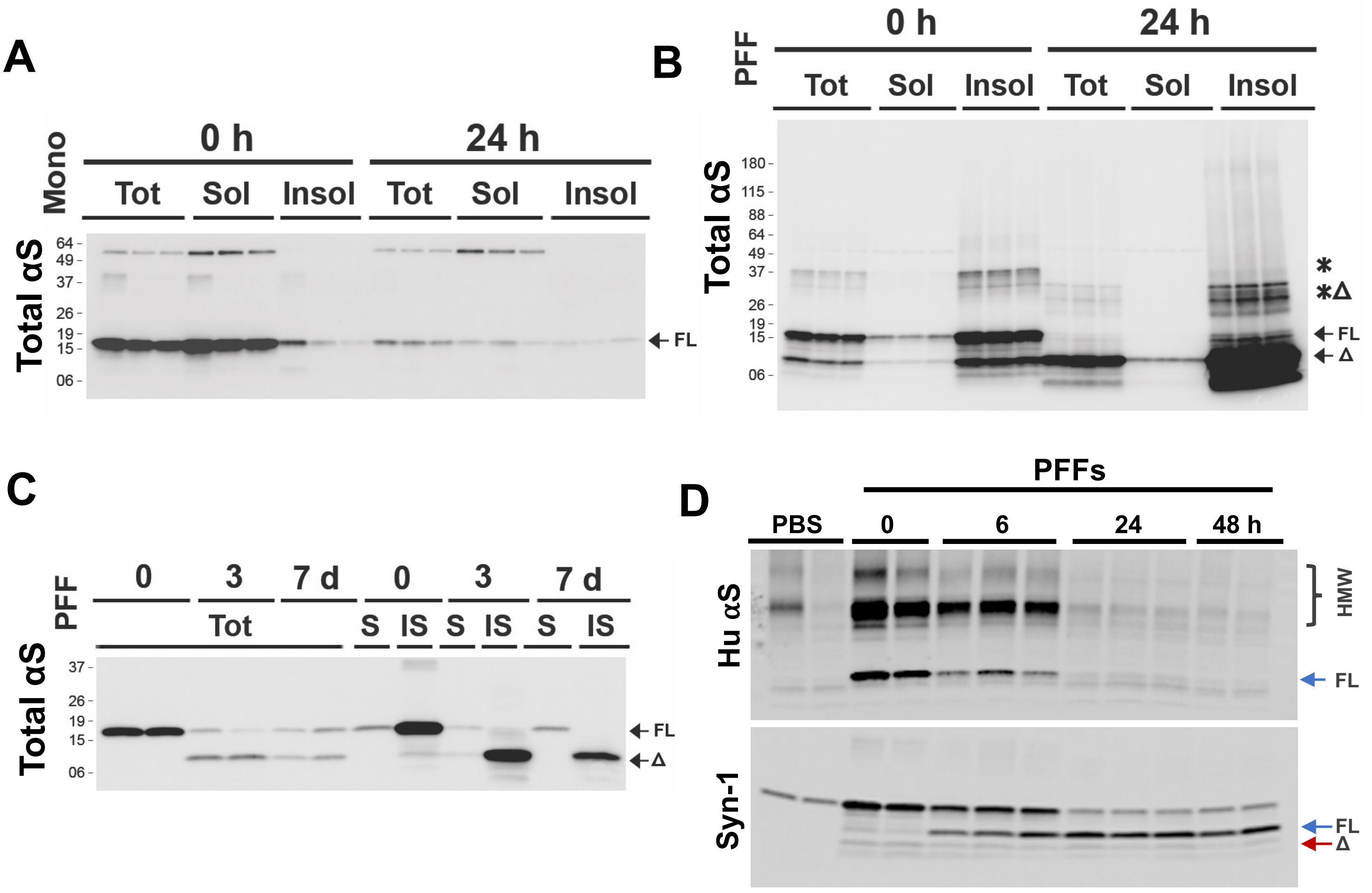


**Figure S3. Internalized αS PFF and truncated αS remain detergent-insoluble. A, B)** Neuronally differentiated CLU198 cells were treated with 4 ug/ml of αS monomers (**A**) or αS PFF **(B)**. At 0h and 24h after washing, the cell lysates were fractionated into total SDS-soluble lysates (Tot), Triton X-100 (TX-100) soluble (Sol) fraction, and TX-100 insoluble (Insol) fraction. Immunoblot analysis show that internalized αS monomers partition with the TX-100 Sol fraction **(A)** while bulk of αS PFF partitions with the TX-100 Insol fraction **(B)**. **C)** PCN treated with PFFs were analyzed for TotαS in TX-100 soluble (S) and insoluble (IS) fractions at 3- and 7-days post αS PFF treatment. Even at 7 days following the initial internalization of αS PFF, virtually all of the truncated αS (Δ) partitions with the insoluble fraction. **D) Metabolism of the C-terminal αS epitope in neurons.** Primary cortical neurons (PCN) were pre-incubated for 2h with 4 μg/ml αS PFF. After washing off excess PFF, cells were replenished with full media and incubated for the indicated time. Total lysates were used for αS Immunoblot analysis using HuαS specific (top) and Syn-1 (bottom) antibodies. Since the epitope for HuαS antibody is located within the C-terminal region (amino acids 115-122), the results show that internalized αS is C-terminally truncated.

**Figure S4.**

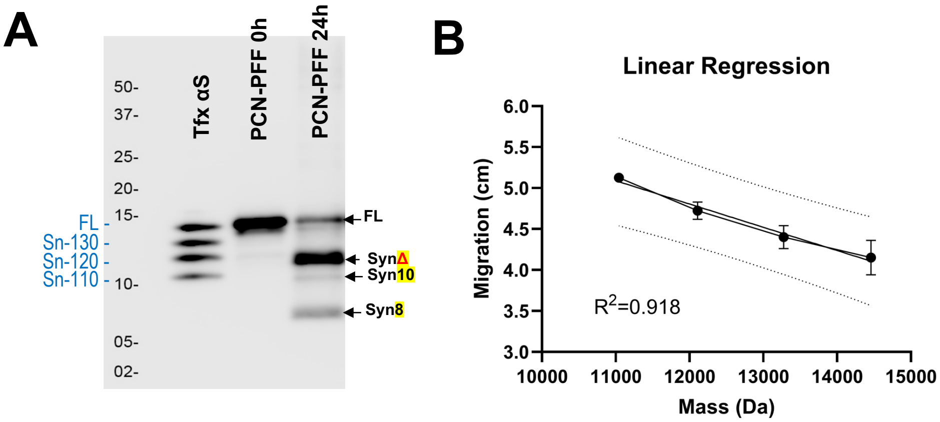


| Hu αS Species (Source) | Predicted Mol Mass, Da | W. Blot Migration (cm) | |
|---|---|---|---|
| Sn-110 (HEK) | 11044 | 5.15 | 5.10 |
| Sn-120 (HEK) | 12111 | 4.80 | 4.65 |
| Sn-130 (HEK) | 13277 | 4.50 | 4.30 |
| Sn-Full (HEK) | 14460 | 4.30 | 4.00 |
| Major Tr-αS (PCN) | 11631 | **4.91** | |

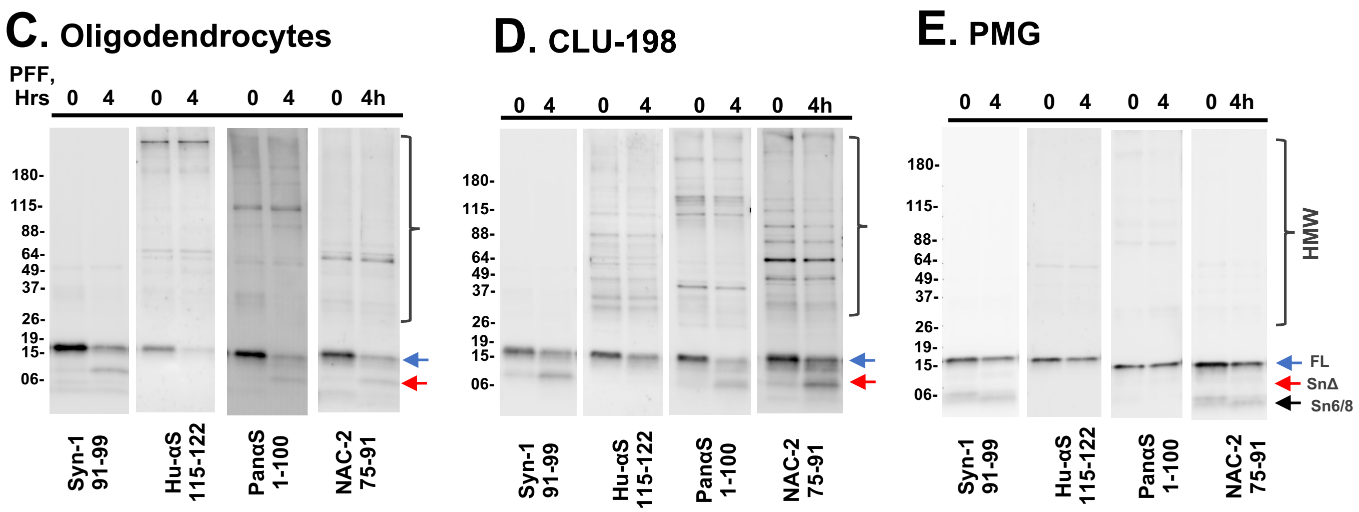


**Figure S4. A, B) Molecular mass estimation of the truncated αS from αS PFF in the primary cortical neurons (PCN).** Total lysates from αS PFF treated PCN, along with cell lysates expressing full-length (FL) and C-terminally truncated HuαS variants, were resolved on a Tris-Tricine gels and immunoblotted for total αS. Based on the relative migration of the transfected αS variants (B), the major truncated αS generated from αS PFF resolves at ~11.5 kDa. **C-E) Epitope mapping of internalized αS epitope in neuronal and glial cells.** Primary oligodendrocytes **(C),** neuronally differentiated CLU-198 cell **(D)**, and Primary microglia (PMG) **(E)** were treated with 4 µg/ml αS PFF and total lysates were used for epitope mapping of the truncated αS. In all cells, the C-terminal HuαS epitope is missing in the truncated variants. While the N-terminal epitope recognized by the Pan-αS antibody is present in FL and Sn10 variants, the N-terminal region is missing in lower MW variant(s) (e.g.Sn6).

Fig. S5

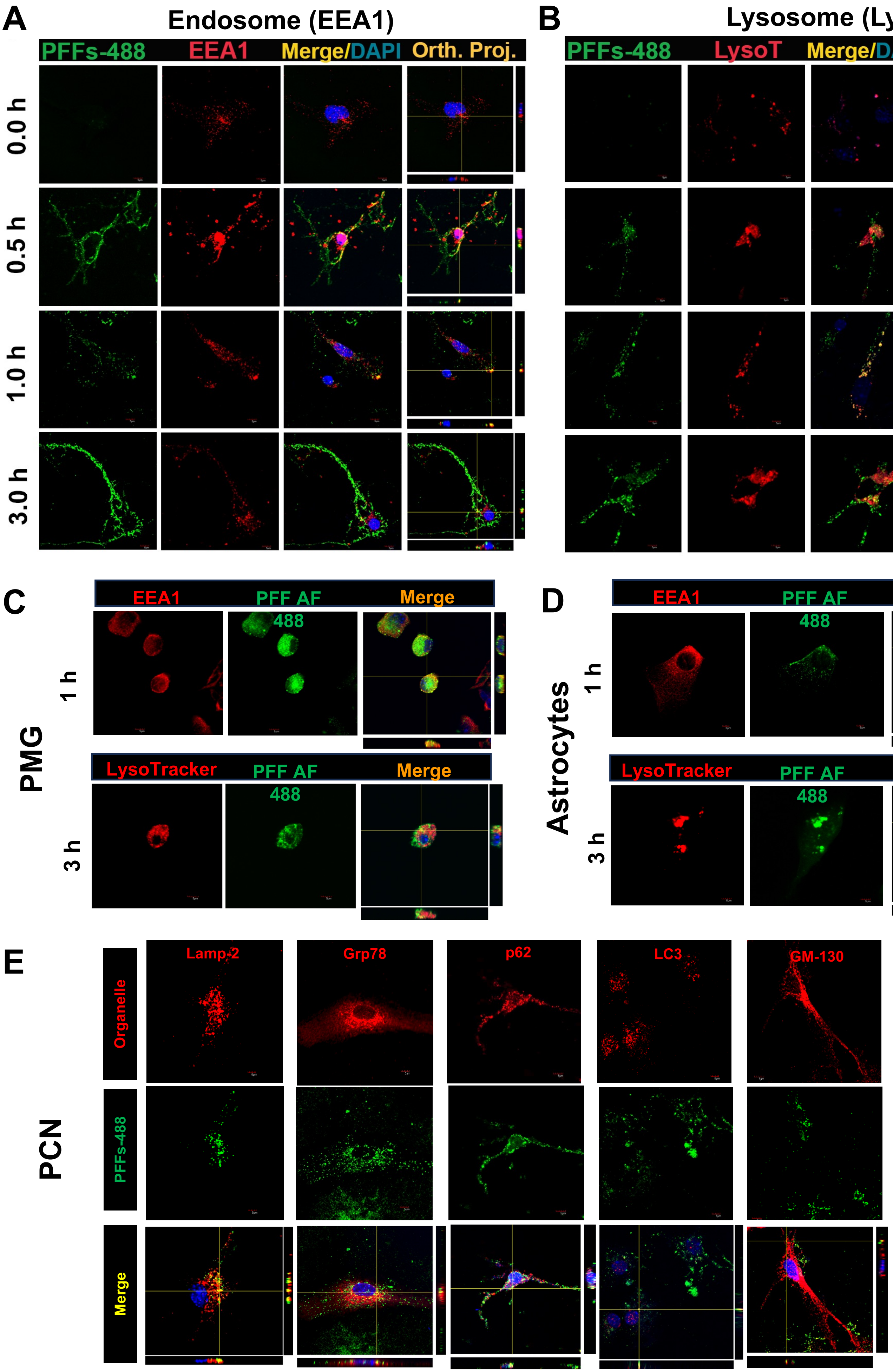
A
Endosome (EEA1)
PFFs-488
EEA1
Merge/DAPI
Orth. Proj.
0.0 h
0.5 h
1.0 h
3.0 h
B
Lysosome (LysoT)
PFFs-488
LysoT
Merge/DAPI
Orth. Proj.
C
EEA1
PFF AF 488
Merge
1 h
PMG
LysoTracker
PFF AF 488
Merge
3 h
D
EEA1
PFF AF 488
Merge
1 h
Astrocytes
LysoTracker
PFF AF 488
Merge
3 h
E
Lamp-2
Grp78
p62
LC3
GM-130
Organelle
PFFs-488
Merge
PCN

**Figure S5. Cellular localization of exogenous αS PFF in cultured cells. A, B)** Individual panels associated with merged images shown in **Figs. 5A** and **5D**. PCN was treated with PFF-AF488 and fixed at 0-, 0.5-, 1- and 3-h following PFF-AF488 addition. Cells were immunostained for early endosome marker EEA1 **(A)** or Lysotracker Red (LysoT) **(B). C,D) Internalized αS PFFs traffics to endosome and Lysosomes in glial cells.** Primary microglia (PMG) **(B)** and astrocytes **(C)** were treated with PFFs-AF488 and colocalized with the early endosome marker (EEA1) (red) at 1 h and Lysosomes (LysoTracker) at 3 h. **E) Localization of internalized αS pre-formed fibrils (PFFs) with intracellular organelles.** Primary cortical neurons (PCN) were treated with PFFs/PFFs-AF488 for 3 hours and the cells were washed with trypsin to remove excess PFFs before fixing cells. PFFs-AF488 (green) were colocalized with various organelle markers: Lysosome, Lamp-2; Endoplasmic Reticulum, Grp-78; Lysosomal substrate, p62; Autophagosome, LC3; Golgi, GM-130.

### Figure S6.

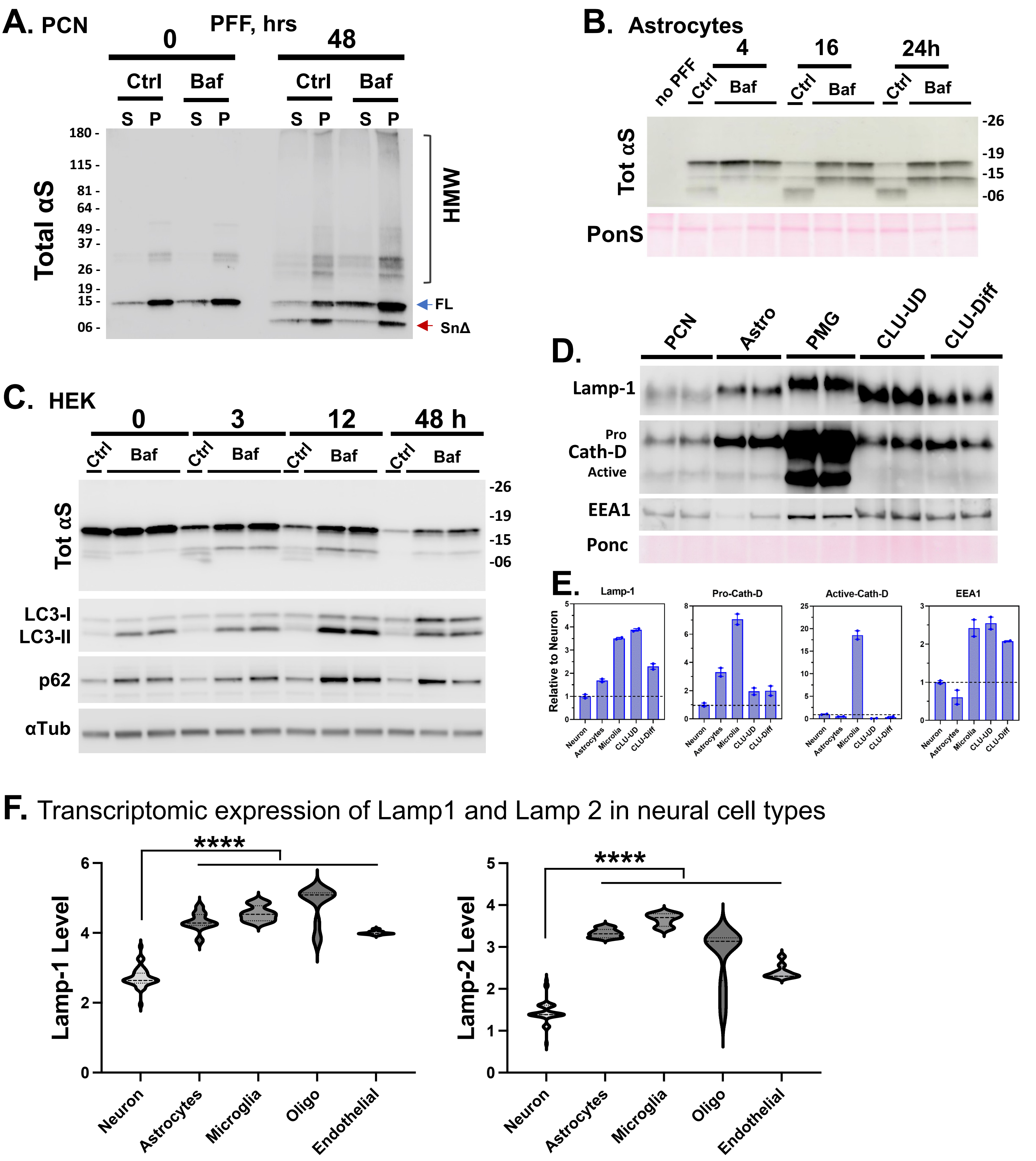
A. PCN
PFF, hrs
0
48
Ctrl
Baf
S P
Total αS
HMW
FL
SnΔ
B. Astrocytes
no PFF
4
16
24h
Tot αS
PonS
C. HEK
0
3
12
48 h
LC3-I
LC3-II
p62
αTub
D.
PCN
Astro
PMG
CLU-UD
CLU-Diff
Lamp-1
Pro
Cath-D
Active
EEA1
Ponc
E.
Lamp-1
Pro-Cath-D
Active-Cath-D
EEA1
Relative to Neuron
Neuron
Astrocytes
Microglia
CLU-UD
CLU-Diff
F. Transcriptomic expression of Lamp1 and Lamp 2 in neural cell types
****
Lamp-1 Level
Lamp-2 Level
Neuron
Astrocytes
Microglia
Oligo
Endothelial

**Figure S6. A) Lysosomal inhibition stabilizes both low and high molecular weight αS.** Mouse primary cortical neurons (PCN) were pre-incubated for 2h with 100 ηM bafilomycin A1 prior addition of 4 μg/ml pre-formed fibrils αS (PFF) for 2h. After trypsin-wash, cells were replenished with full media and Baf for the indicated time. Triton X-100 soluble (S) and insoluble (P) fractions were separated and analyzed for Tot αS (Syn-1) and Pan-S by immunoblotting. **B, C) Lysosome inhibition stabilizes internalized αS PFF in astrocytes (B) and HEK293 cells (C).** Cells were pre-incubated for 100 nM Baf and 4 μg/ml αS-PFF and lysates were collected at indicated times. Immunoblot analysis of αS shows that while most of αS is metabolized by 12-16 hours in astrocytes (**B**) and HEK293 cells (**C**). Baf treatment leads to continued accumulation of αS, even at 24-48 h. Also shown is Ponceau-S total protein stain or α-tubulin immunoblot to control for equal loading. Immunoblot analysis of LC3-I/-II and p62 verify the inhibition of lysosomes. **D, E) Neuronal cells have lower lysosomal markers. D)** Primary cortical neuron (PCN), primary astrocytes, primary microglia (PMG), and differentiated or undifferentiated mouse hippocampal CLU cells were harvested, and lysosomal markers (Lamp-1, Cathepsin D, and EEA1) were evaluated by immunoblotting. **E)** Quantitative analysis of immunoblot shown in D. While statistical analysis was not done, the duplicate samples clearly show that lysosomal markers are lower in neurons than in non-neuronal cells. **F)** Whole brain cell type expression levels were extracted for Neurons, Astrocytes, Microglia, Oligodendrocytes, and Endothelial cells from DropViz data single cell transcriptomic data base for Lamp1 (http://dropviz.org/?_state_id_=c8c0e49d46a99f4b) and Lamp2 (http://dropviz.org/?_state_id_=89519e3741f26c75). The violin plot of the values show that Neurons express less Lamp1 and Lamp2 than other brain cell types. ****$p<0.0001$*, One Way ANOVA.*

#### Figure S7

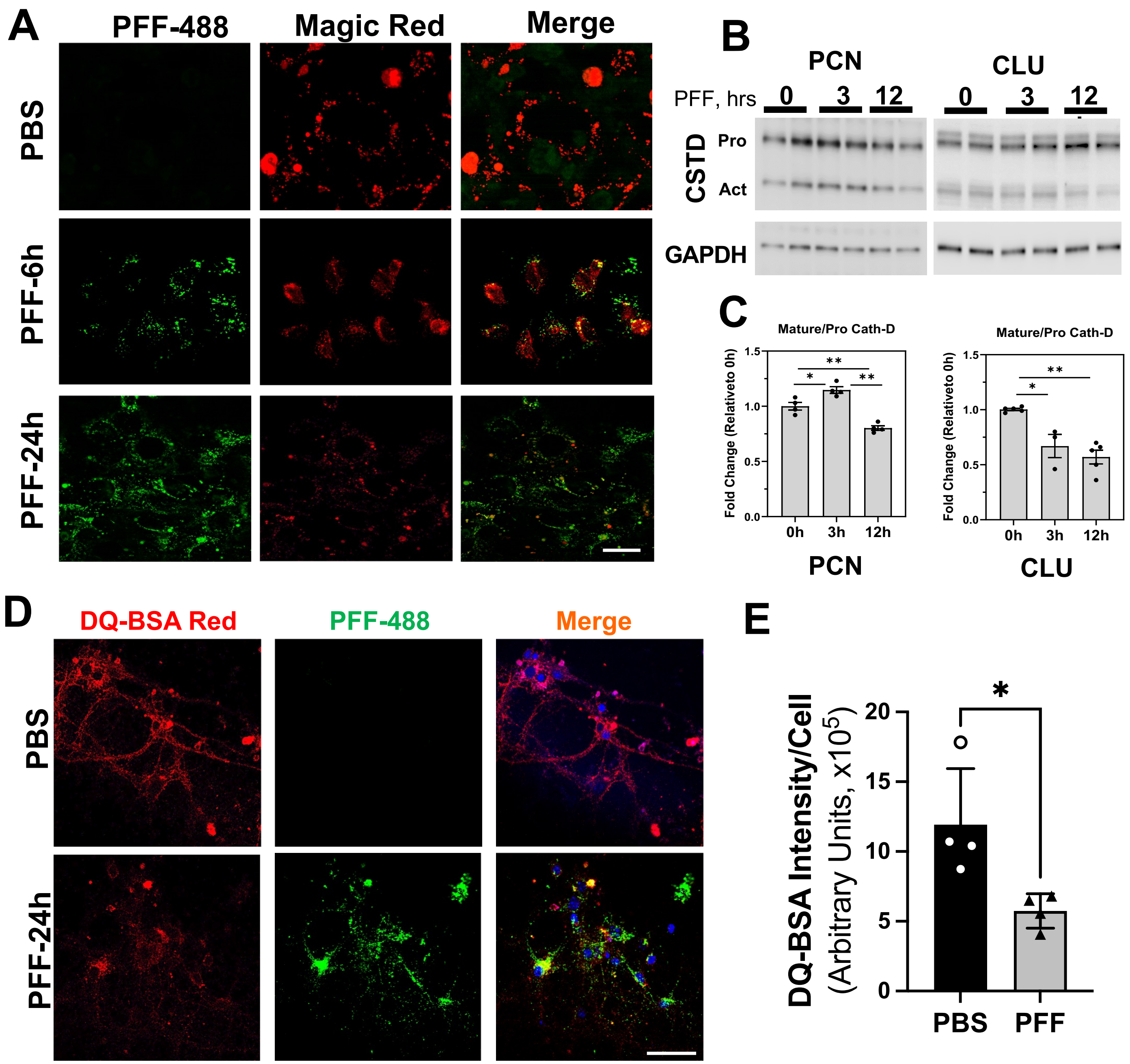


**Figure S7. Internalized αS PFFs inhibit lysosomal function in neuronal cells. A)** Differentiated CLU198 cells were treated with PFF-AF488, washed, and imaged at 6- or 24-h for PFF-AF488 or Magic Red. The Magic Red signal appears reduced in PFF-treated cells. Quantitative analyses of 24 h time point is shown in Fig. 7. Bar=10 µm. **B)** PFF treated PCN or CLU198 cells were analyzed for the relative amount of active and pro Cathepsin D (CSTD) by immunoblotting. **C)** Immunoblot shown in B were quantified showing that the PFF treatment reduces relative levels of active Cath-D, relative to the pro-Cath-D. *$p<0.05$, **$p<0.01$, One-Way Anova, Tukey's multiple comparison test. **D, E)** Primary cortical neurons were treated with PFF-AF488, washed, and incubated for 24-h and treated with DQ-BSA Red, which is hydrolyzed in functional lysosomes to produce red fluorescence. Confocal images show that PFF treatment is associated with reduced DQ-BSA Red signal **(D)**. Quantitative analysis of total DQ-BSA Red signal intensity, normalized to the number of cells, confirms that PFF treatment leads to lysosomal dysfunction in neurons **(E)**. *$p<0.05$, unpaired t-test, n=4 independent cultures. Bar=50 µm (C). *$p<0.05$, unpaired t-test.

**Figure S8.**

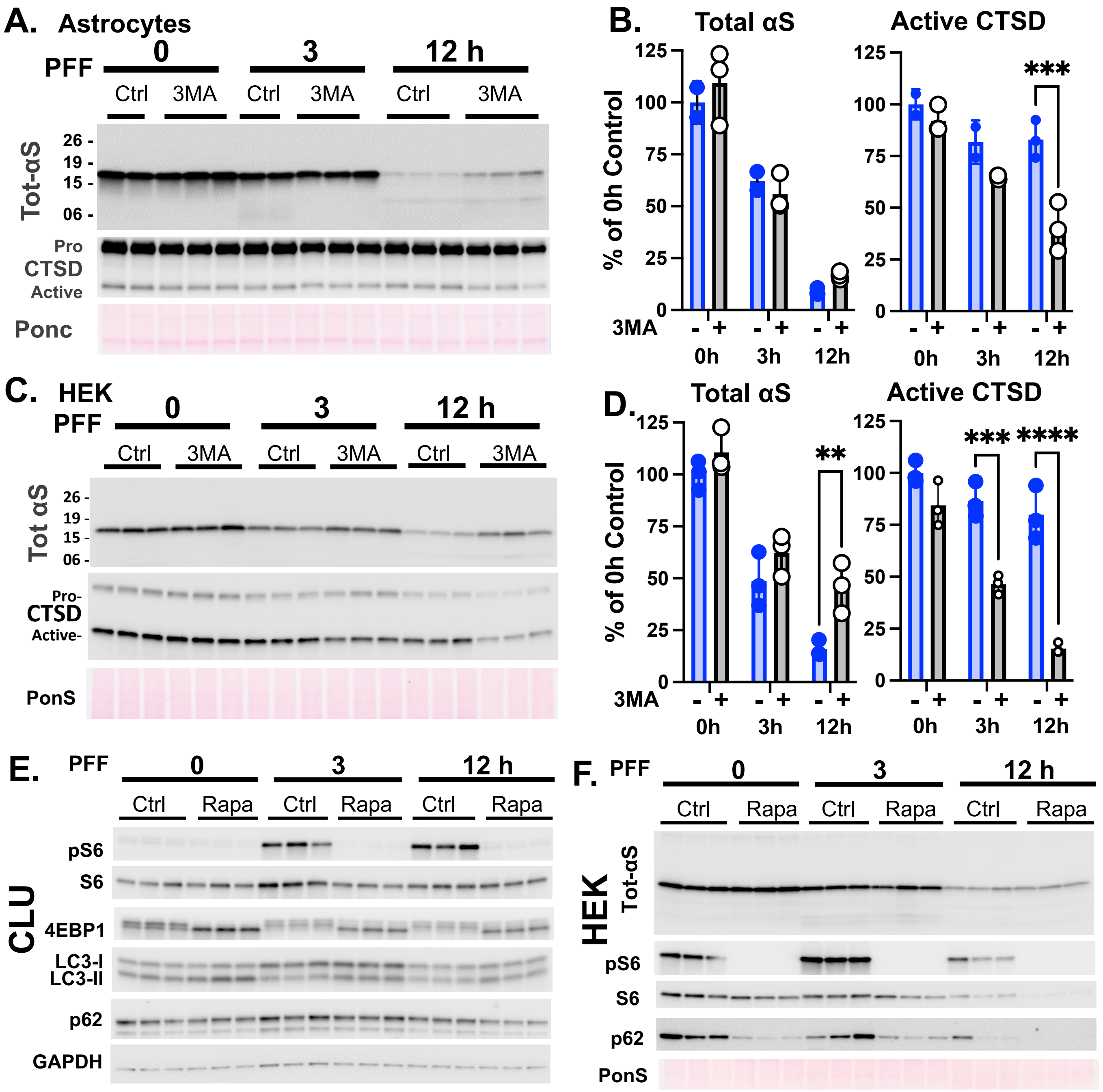


**Figure S8. (A-D) Inhibition of autophagy leads to modest lysosomal inhibition with a modest impact on αS accumulation in astrocytes and HEK293 cells.** Primary cultures of astrocytes (**A, B**) and HEK293 cell line (**C, D**) treated with 3MA to inhibit autophagy as described in Figure 8. The levels of internalized αS PFF and the lysosomal protease, Cathepsin-D (CTSD), determined by immunoblot analysis, show that 3MA treatment leads to a modest but significant increase in αS levels in HEK293 cells (**C, D**) and decreased active CTSD levels in both Astrocytes and HEK293 cells (**B, D**). ***p<0.01, ***p<0.001, ****p<0.0001*, Two-way ANOVA. **(E, F) Rapamycin increases autophagy but does not affect αS PFF metabolism. (E**) CLU-198 shown in Fig. 8D analyzed for mTOR inhibition (pS6, 4EBP) and autophagy (LC3-II and p62). In addition to inhibition of mTOR, Rapa treatment leads to increased LC3-II and reduced p62, indicating to enhanced autophagy. **(F)** In HEK293 cells, Rapa treatment does not affect the metabolism of internalized αS.

**Figure S8.**

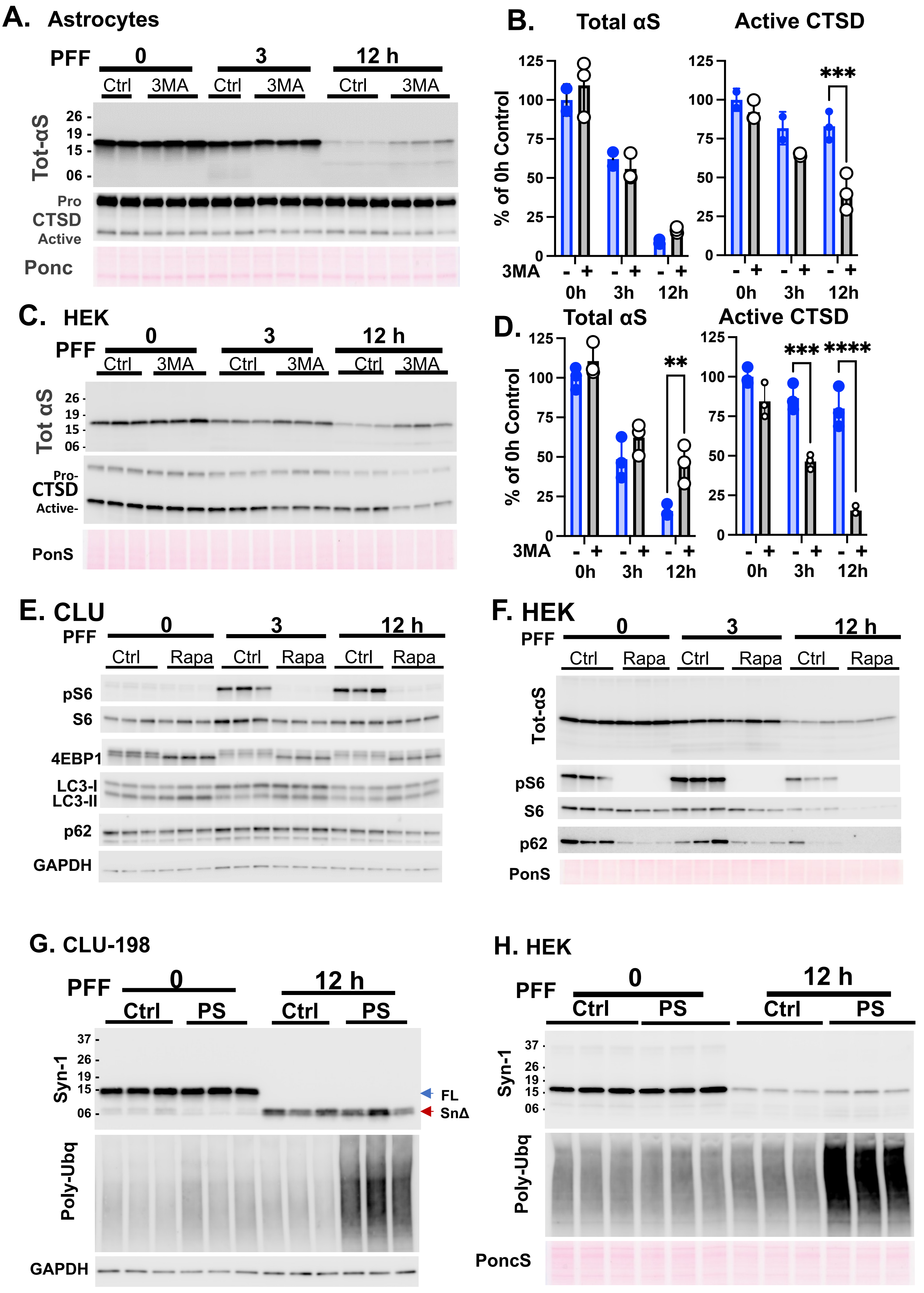

A. Astrocytes
PFF 0 3 12 h
Ctrl 3MA Ctrl 3MA Ctrl 3MA
Tot-αS
26 - 19 - 15 - 06 -
Pro CTSD Active
Ponc
B. Total αS
Active CTSD
% of 0h Control
***
3MA - + - + - +
0h 3h 12h
C. HEK
PFF 0 3 12 h
Ctrl 3MA Ctrl 3MA Ctrl 3MA
Tot αS
Pro- CTSD Active-
PonS
D. Total αS
Active CTSD
**
*** ****
E. CLU
PFF 0 3 12 h
Ctrl Rapa Ctrl Rapa Ctrl Rapa
pS6
S6
4EBP1
LC3-I LC3-II
p62
GAPDH
F. HEK
Tot-αS
PonS
G. CLU-198
PFF 0 12 h
Ctrl PS Ctrl PS
Syn-1
37 - 26 - 19 - 15 - 06 -
FL
SnΔ
Poly-Ubq
GAPDH
H. HEK
PoncS

**Figure S8. A-D) Inhibition of autophagy leads to modest lysosomal inhibition with a modest impact on αS accumulation in astrocytes and HEK293 cells.** Primary cultures of astrocytes (**A, B**) and HEK293 cell line (**C, D**) treated with 3MA to inhibit autophagy as described in Figure 8. The levels of internalized αS PFF and the lysosomal protease, Cathepsin-D (CTSD), determined by immunoblot analysis, show that 3MA treatment leads to a modest but significant increase in αS levels in HEK293 cells (**C, D**) and decreased active CTSD levels in both Astrocytes and HEK293 cells (**B, D**). ***p<0.01, ***p<0.001, ****p<0.0001*, Two-way ANOVA. **E, F) Rapamycin increases autophagy but does not affect αS PFF metabolism.** (**E**) CLU-198 shown in Fig. 8D analyzed for mTOR inhibition (pS6, 4EBP) and autophagy (LC3-II and p62). In addition to inhibition of mTOR, Rapa treatment leads to increased LC3-II and reduced p62, indicating to enhanced autophagy. **(F)** In HEK293 cells, Rapa treatment does not affect the metabolism of internalized αS. **G, H) Proteosome inhibition does not impact αS PFF processing/metabolism.** (**G**) Differentiated mouse hippocampal cell CLU-198 and **(H)** Human embryonic kidney HEK cells were pre-incubated with 15 nM PS-341 (PS; 4h) and 4 µg/ml pre-formed fibrils αS (PFFs) for 2h before the trypsin-wash. After the trypsin wash, cells were replenished with full media and PS for the indicated time. Tot αS (Syn-1), Poly-Ubiquitin (Poly-Ubq), and GAPDH were detected by immunoblotting analysis in G. Ponceau S was used in **H**. PS-341 treatment do not impact αS truncation or metabolism in CLU-198 cells (**G**) and does not impact αS metabolism in HEK293 cells (**H**).

### Figure S9.

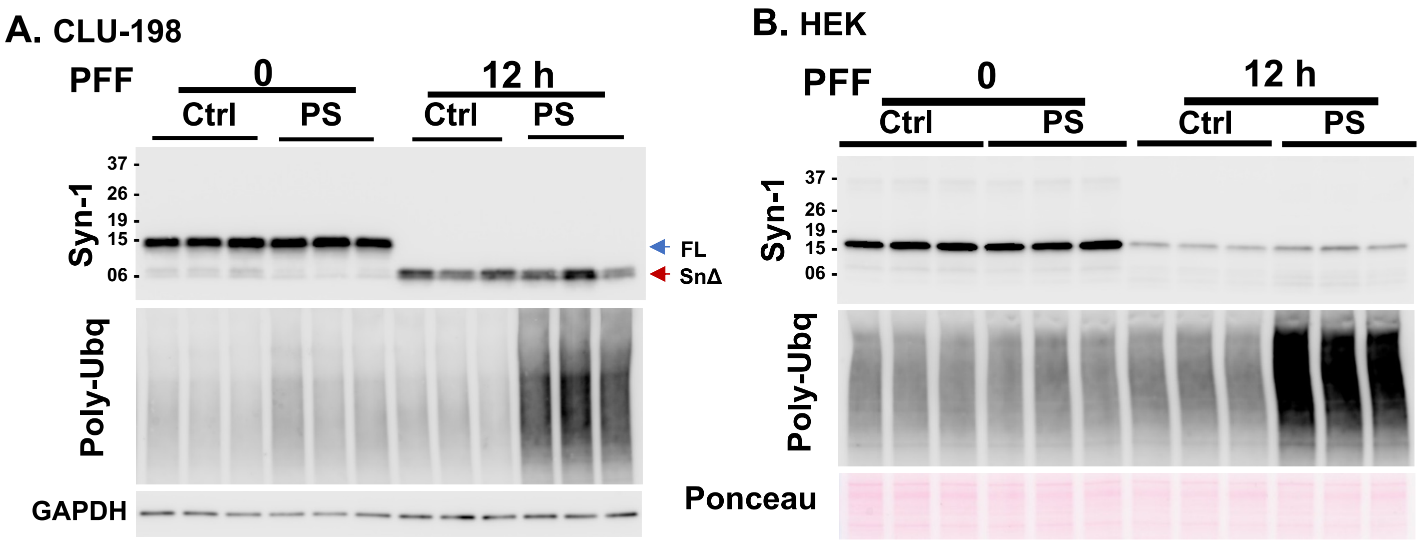


**Figure S9. Proteosome inhibition does not impact αS PFF processing/metabolism. (A)** Differentiated mouse hippocampal cell CLU-198 and **(B)** Human embryonic kidney HEK cells were pre-incubated with 15 nM PS-341 (PS; 4h) and 4 µg/ml pre-formed fibrils αS (PFFs) for 2h before the trypsin-wash. After the trypsin wash, cells were replenished with full media and PS for the indicated time. Tot αS (Syn-1), Poly-Ubiquitin (Poly-Ubq), and GAPDH were detected by immunoblotting analysis. Ponceau S was used in **B**. PS-341 treatment do not impact αS truncation in CLU-198 cells (A) and does not impact αS metabolism in HEK293 cells (**B**).
